# Vascularized Human Cardiac Organoids Reveal Endothelial–Cardiomyocyte Crosstalk and Mechanisms of Carfilzomib-Induced Cardiotoxicity

**DOI:** 10.64898/2026.04.14.718448

**Authors:** Yuan Yang, Liyang Zhao, Honglin Tao, Yao Zhang, Hanwen Wang, Henghui Xu, Ning Ma, Ning-Yi Shao, Joe Z. Zhang

## Abstract

Human cardiac organoids (hCOs) provide a powerful platform for modeling heart development and disease, yet their lack of vasculature limits physiological relevance. Here, we generate vascularized human cardiac organoids (vhCOs) by assembling hCOs with blood vessel organoids, enabling the self-organized formation of unified tissues with extensive vascular networks and substantially enhanced cardiomyocyte (CM) maturation. Single-nucleus multiomic profiling reveals CM state resembling the 17-week human fetal heart, while endothelial cells (ECs) acquire cardiac-specific and arterial-like identities. Integrative cell–cell communication analyses and experimental validation identify CM-derived Laminin-α2 as a driver of arterial EC specification, while reciprocal *EFNB2* signaling from arterial ECs promotes CM maturation, uncovering bidirectional cardiac–vascular crosstalk. Modeling carfilzomib-induced cardiotoxicity in vhCOs reveals ATF4-dependent endoplasmic reticulum (ER) stress and IL8-mediated inflammatory responses that are blunted in non-vascularized hCOs. Pharmacological attenuation of ER stress alleviates carfilzomib-induced cardiotoxicity. Altogether, vhCOs provide a developmentally advanced and physiologically faithful human cardiac model.

## INTRODUCTION

The human heart comprises multiple cell types, including cardiomyocytes (CMs), endothelial cells (ECs), fibroblasts (FBs), and others, which coordinate to maintain contractile function and tissue homeostasis. However, studying human heart development and cardiovascular disease has long been challenging due to the paucity of available human heart tissue. Human cardiac organoids (hCOs) derived from pluripotent stem cells (PSCs), including induced pluripotent stem cells (iPSCs) and embryonic stem cells (ESCs), have recently emerged as powerful platforms for studying human fetal heart development^1–8^ and a wide spectrum of cardiovascular diseases.^9–15^

Despite the significant advances offered by hCOs, a major unresolved limitation is their poor vascularization. The absence of a functional vascular system limits nutrient and oxygen diffusion, an issue that becomes particularly pronounced as organoids grow to millimeter scale during long-term culture, often resulting in substantial apoptosis or necrosis at the organoid core.^15,16^ This shortcoming severely restricts organoid size, viability, and functional maturation, limiting their potential for modeling later developmental stages and precise disease modeling. Beyond perfusion, the vascular niche via direct cell-cell contacts and paracrine signaling has been shown to play a critical role in regulating CM maturation, structural organization, and electrophysiological function.^17–20^

To address these limitations, recent efforts have been made to incorporate vascular cells into hCOs using bioengineering approaches or microfluidic “organ-on-chip” systems. Co-culture of primary or PSC-derived ECs promotes the formation of EC-enriched networks within cardiac microtissues or organoids.^19,21–24^ Alternatively, the integration of microfluidic platforms has enabled the formation of perfusable vascular system in “heart-on-chip” models.^24–27^ However, most of these studies typically rely on predefined mixtures of separately differentiated cardiovascular cells, resulting in limited cellular diversity and restricted ability of ECs to form vessel-like structure. More recently, micropattern-based strategies have been used to generate vascularized hCOs and these organoids resemble a 6.5-week human fetal heart in both structure and function.^20^ Yet, the geometric confinements imposed by micropatterns restrict the natural self-organization and may limit physiological cell-cell interactions. Moreover, previous work has primarily emphasized how vascularization modulates CM maturation, while the reciprocal influence of the cardiac microenvironment on endothelial identity and vascular niche specification remains largely unknown.

Here, we establish vascularized human cardiac organoids (vhCOs) by integrating hCOs with human blood vessel organoids (hBVOs) within a matrix. This assembly strategy enables the efficient generation of vhCOs containing diverse cardiac and non-cardiac cells in proportions comparable to those of the human fetal heart at 17-post-conceptional weeks (PCWs). The incorporation of functional vasculature-like networks not only markedly reduces apoptosis at the organoid core, but also significantly enhances the functional and metabolic maturation of CMs within vhCOs similar to that of 17-PCW human fetal heart mediated by Ephrin-B2–EphA4 signaling pathway. Moreover, compared with ECs in hBVOs, the cardiac microenvironment within vhCOs promotes a cardiac-specific endothelial lineage identity including enhanced differentiation toward arterial subtypes as regulated by Laminin-α2, thereby achieving greater physiological resemblance to the human heart.

Building on vhCOs with cardiac-specific vasculature and advanced maturation, we next investigated the cardiovascular toxicity of Carfilzomib (CFZ), an irreversible proteasome inhibitor approved for the treatment of refractory/relapsed multiple myeloma (MM). Accumulating evidence indicates that CFZ can cause heart failure, arrhythmias, and hypertension, implicating detrimental effects on multiple cardiovascular cell types.^28–35^ Several studies have demonstrated the distinct drug responses between isolated CM cultures and multicellular co-culture systems. ^19,22,27,36,37^ Moreover, traditional 2D cultures and non-vascularized hCOs lack the tissue microenvironment and functional vasculature, limiting their capacity to fully model CFZ-induced cardiotoxicity. Using vhCOs, we successfully recapitulated the CFZ-induced cardiac dysfunction accompanied by augmented endoplasmic reticulum (ER) stress response, leading to increased secretion of Interleukin-8 (IL8) in CFZ-treated vhCOs, but not in hCOs or hBVOs. Single-nucleus multiomic analysis further corroborated these findings by showing the positive correlation of expression between ER stress-associated genes such as *ATF4* and *IL8* in CFZ-treated vhCOs. We finally identified that the chemical chaperone 4-phenylbutyric acid (4-PBA) can rescue CFZ-induced cardiotoxicity. Altogether, these findings demonstrate the potential of vhCOs as an advanced model for dissecting disease mechanisms and cardioprotective drug discovery.

## RESULTS

### Generation and characterization of vhCOs

To assemble vhCOs by integrating functional hCOs with hBVOs *in vitro* (Figure 1A), we first generated hCOs from hiPSCs using a stepwise differentiation protocol with small molecules and growth factors (Figure S1A).^2,3^ This approach efficiently generated spontaneously beating hCOs across multiple hiPSC lines with high reproducibility (Figure S1B). During differentiation, cardiac marker genes were progressively upregulated, whereas the expression levels of endothelial marker genes remained low, mirrored by the minimal vasculature observed by whole-mount immunostaining of hCOs (Figures S1C and S1D). Single-nucleus multiomic profiling (snRNA-seq and snATAC-seq) of hCOs on days 15 and 42 showed that CMs (clusters CM1-4) made up the predominant cell population, while ECs accounted for only 1.3% of total cells (Figures S2A-S2D), a proportion consistent with previous studies, which typically find ECs comprising 1%-3% of the total population.^38^ Together, these data demonstrate that current methods reliably generate CM-rich hCOs with minimal ECs and vasculature-like structures.

**Figure 1.**
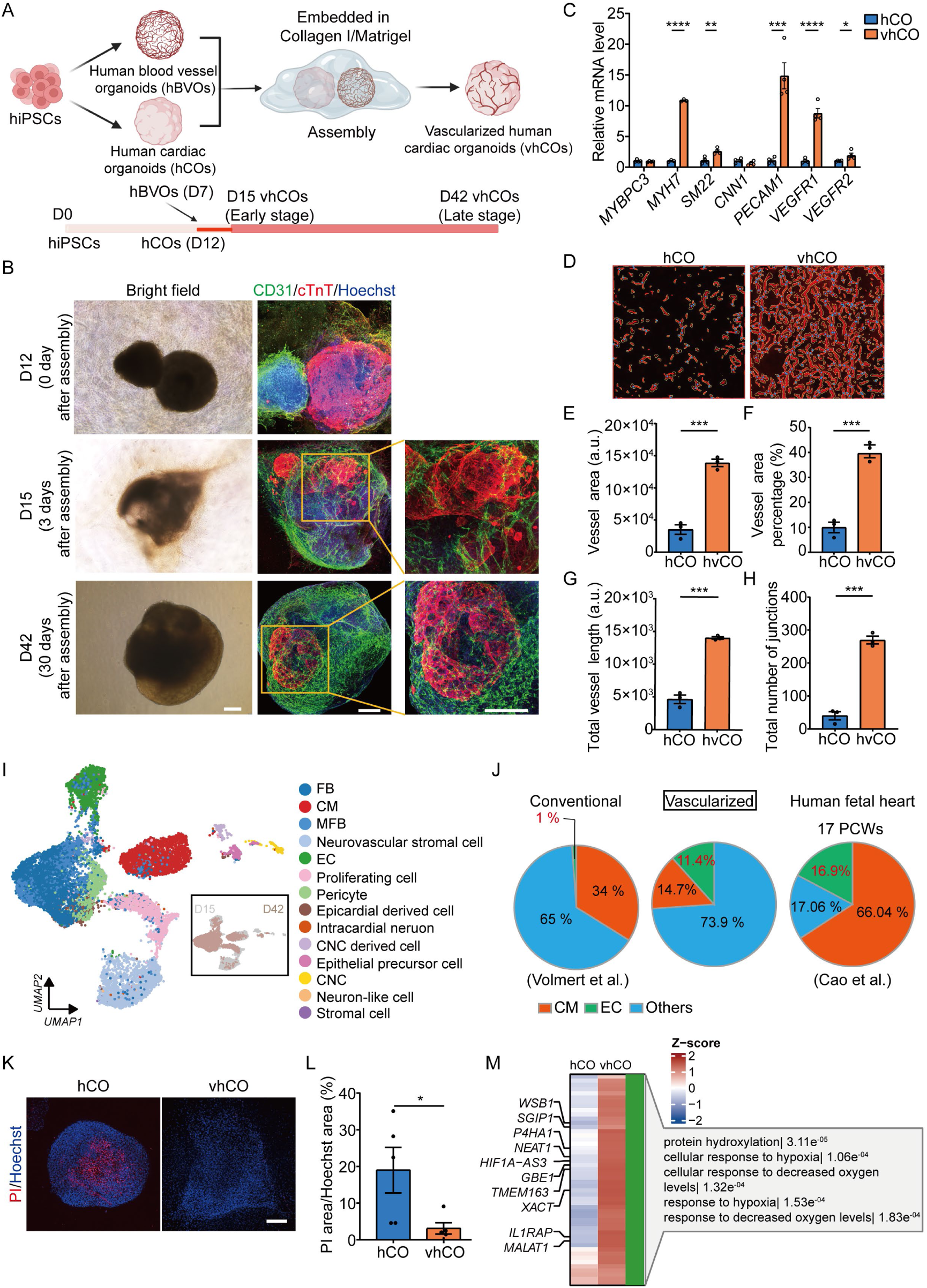
Generation and characterization of vhCOs. **(A)** Schematic diagram of the assembly of hCOs with hBVOs to generate vhCOs. **(B)** Representative bright-field and whole-mount immunostaining showing CD31 and cardiac troponin T (cTnT) expression in vhCOs after 0-, 3-, and 30-day assembly. **(C)** Temporal expression of CM, EC, and mural cell marker genes in D42 hCOs and vhCOs. (n = 3 biological replicates). **(D)** Representative binary images of CD31^+^ vasculature. **(E-H)** Quantification of vascular network parameters, including total vessel area (E), vessel area percentage (F), total vessel length (G), and total number of junctions (H) analyzed from CD31-stained images using AngioTool. (n = 3 biological replicates). **(I)** UMAP visualization of integrated snRNA-seq data from D15 and D42 vhCOs, identifying 14 transcriptionally distinct clusters. Major cell types are annotated. Inset shows D15 (brown) and D42 (gray) cells. **(J)** Comparison of cellular composition among conventional hCOs,^38^ vhCOs, and 17-PCW human fetal heart tissue^41^ based on snRNA-seq. **(K)** Representative fluorescence images of propidium iodide (PI; red) and Hoechst (blue) staining in D42 hCOs and vhCOs. (n = 5 biological replicates). **(L)** Quantification of PI^+^ area normalized to total Hoechst^+^ area. **(M)** Heatmap of differentially expressed genes between hCOs and vhCOs. Scale bars, 200 μm. Data are presented as mean ± SEM. Statistical significance was determined by unpaired two-tailed Student’s *t* test. *p<0.05, **p<0.01, ***p<0.001, ****p<0.0001.

To overcome this limitation, we generated hBVOs from hiPSCs by stepwise induction of mesoderm, endothelial lineages, and vascular sprouting (Figure S3A).^39^ During differentiation, expression of endothelial and mural cell genes significantly increased (Figure S3B). Whole-mount immunostaining of day-12 hBVOs showed CD31⁺ vessel-like structures surrounded by PDGFRβ⁺ pericytes and vimentin⁺ mural cells, indicative of a multilayered vascular structure (Figure S3C). When transplanted under the kidney capsule of NSG mice, Texas red-dextran perfusion revealed anastomosis between hBVO-derived vasculature and host blood vessels (Figures S3D and S3E), demonstrating functionally perfusable vascular networks. Isolated ECs from hBVOs efficiently internalized DiI-acetylated low-density lipoprotein (DiI-Ac-LDL), confirming endothelial activity (Figures S3D and S3F). Single-nucleus multiomic profiling identified five major cell clusters: ECs, mural cells, neural-like cells, proliferating cells, and neurovascular stromal cells with distinct chromatin accessibility landscapes (Figures S4A-S4D), indicating a mature and diverse vascular niche. Together, these findings confirm that hBVOs form structurally complete and functionally competent vessel organoids.

We next sought to generate vhCOs by integrating hCOs and hBVOs into a unified organoid. Since embedding day-7 hBVOs into matrix is essential for their development, we assembled day-12 hCOs with day-7 hBVOs in a Collagen I-Matrigel matrix. Optimization of assembly conditions identified a 1:1 ratio of hCO:hBVO and a 1:1 mixture of chemically defined medium (CDM) and RPMI as the optimal condition (Figure 1B and Figures S5A-S5E). Initially, these two organoids remained juxtaposed, with CD31⁺ vessels restricted to the hBVO (Figure 1B). By day 15 (3 days post-assembly), CD31⁺ vascular networks extended across the contact interface and began infiltrating the cTnT⁺ cardiac region. By day 42, these two organoids had assembled into a single organoid, with branched and lumenized CD31⁺ vascular plexus enveloping and penetrating the cardiac tissue (Figures S5F and S5G, and Supplementary Video S1), indicative of progressive vascular ingrowth and remodeling. qPCR analysis further confirmed successful vascular integration, showing increased expression of the mural cell markers (*SM22*) and key angiogenic genes (*PECAM1*, *VEGFR1*, and *VEGFR2*) in vhCOs compared with hCOs (Figure 1C). Notably, the CM maturation marker *MYH7*, predominantly expressed in adult CMs,^40^ was markedly upregulated in vhCOs, suggesting that the introduced vascular niche enhances CM maturation. Moreover, hBVO-derived vascular networks in vhCOs exhibited increased vessel area, length, and branching compared with those in hCOs (Figures 1D-H).

Single-nucleus multiomic profiling of vhCOs on days 15 and 42 identified 14 cell clusters corresponding to expected cardiac and vascular cell types, including CMs, FBs, ECs, proliferating cells, and pericytes (Figure 1I). ECs accounted for 11.4% of total cells, substantially higher than in hCOs (1.3%),^38^ demonstrating successful vascular augmentation (Figure 1I and Figure S2B). Strikingly, the cellular composition of vhCOs closely resembled that of the human fetal heart at 17 PCWs (Figure 1J),^41^ supporting the establishment of a developmentally advanced and diverse cardiac-vascular niche. A major challenge in long-term organoid culture is hypoxia-induced apoptosis in the core of large organoids due to lack of perfusion. Compared with hCOs, vhCOs showed markedly fewer apoptotic cells (Figures 1K-1L). Correspondingly, transcriptomic analysis revealed upregulation of genes responsive to reduced oxygen levels (Figure 1M).

Collectively, these data demonstrate the robust generation of vhCOs featuring integrated and functional vascular networks that support cell survival, enhance tissue maturation, and more closely recapitulate the cellular complexity of the later stage of human developing heart.

### Vascularization promotes CM maturation in vhCOs

To assess the maturation status of CM populations in hCOs and vhCOs, we first integrated our datasets with previously published hiPSC-CM datasets spanning different differentiation stages^42^ and performed Monocle trajectory analysis. UMAP visualization revealed that CMs from hCOs or vhCOs formed two distinct clusters, with vhCO-derived CMs closer to late-stage CMs (Figure 2A). We next assessed the differentiation potential of these populations using CytoTRACE, a computational framework where higher scores indicate greater stemness and lower scores reflect advanced maturation.^43^ vhCO-derived CMs had the lowest CytoTRACE score among all groups (Figure 2B), confirming their superior maturation status. To precisely benchmark their developmental stage, we integrated our snRNA-seq data with published human fetal heart datasets.^41,44,45^ The trajectory analysis demonstrated that vhCO-derived CMs aligned closely with human fetal CMs at approximately 17 PCWs (Figure 2C). Consistent with these developmental differences, vhCO-derived CMs showed increased expression of genes associated with electrophysiological maturation, including the potassium channel *KCNQ1*, the L-type calcium channel *CACNA1C*, and the adult sarcomeric isoforms *TNNI3*, compared with age-matched hCO-CMs (Figures 2D and 2E). Accordingly, the *TNNI3/TNNI1* and *MYH7/MYH6* expression ratios were significantly higher in vhCOs (Figures 2F and 2G), indicating a shift toward a more adult-like contractile phenotype. Functionally, vhCOs demonstrated enhanced calcium handling capability. Calcium imaging revealed higher calcium transient amplitudes and shorter peak-to-half-decay time in vhCOs compared with hCOs (Figures 2H–2J), consistent with elevated expression of *RYR2* and *KCNQ1* (Figures 2K and 2L), as well as increased transcription level and chromatin accessibility of calcium handling genes such as *ATP2A2* and *SLC8A1* (Figures S6A and S6B). Meanwhile, the automaticity-related ion channel genes, like *HCN4*, was downregulated in vhCOs compared with hCOs (Figures S6C). Given that *HCN4* is typically expressed in nodal tissue or immature hiPSC-CMs,^46,47^ its downregulation indicates the acquisition of a more mature CM phenotype in vhCOs. In addition, mitochondrial respiration measurements further supported improved metabolic maturation in vhCOs. Seahorse extracellular flux analysis showed significantly higher basal and ATP-linked respiration, increased maximal respiratory capacity, reduced proton leak, and higher spare respiratory capacity in vhCOs compared with hCOs (Figures 2M–2S).

**Figure 2.**
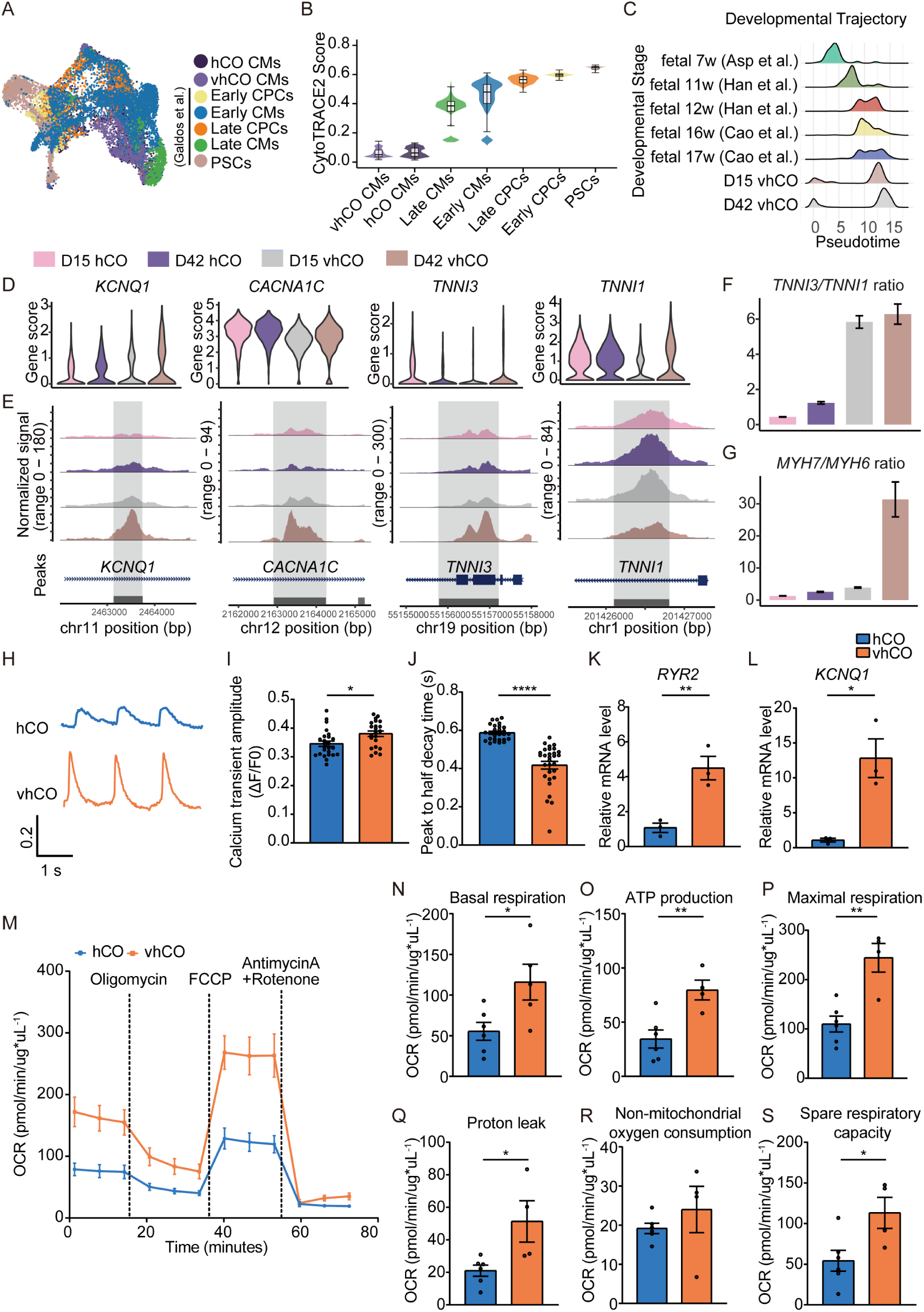
Vascularization promotes transcriptional, functional, and metabolic maturation of CMs in vhCOs. **(A)** UMAP visualization of integrated snRNA-seq data from hCOs, vhCOs, and published datasets from hiPSC-derived pluripotent stem cells (PSCs), early and late cardiac progenitor cells (CPCs), early and late CMs.^42^ **(B)** Comparison of CytoTRACE stemness scores across hiPSC-CM differentiation stages and CMs derived from hCOs and vhCOs. **(C)** Developmental trajectory analysis of hCO- or vhCO-derived CMs relative to human fetal heart CMs across different PCWs.^41,44,45^ **(D-E)** Violin plots (D) and chromatin accessibility tracks (E) showing gene expression and open chromatin at loci encoding maturation-associated ion channel and sarcomeric genes in D42 vhCO-derived CMs compared with D15 vhCO- and age-matched hCO-derived CMs. Genomic positions of peaks are indicated below each locus. **(F-G)** Ratios of *TNNI3/TNNI1* (F) and *MYH7/MYH6* (G) in hCOs and vhCOs. **(H)** Representative calcium transient traces in hCOs and vhCOs. **(I-J)** Quantification of calcium transient amplitude (I) and peak to half-decay time (J). (n = 30-40 biological replicates). **(K-L)** qPCR analysis of *RYR2* (K) and *KCNQ1* (L) expression in hCOs and vhCOs. (n = 3 biological replicates). **(M)** Seahorse extracellular flux analysis of oxygen consumption rate (OCR) in hCOs and vhCOs. **(N-S)** Quantification of basal respiration (N), ATP production (O), maximal respiration (P), proton leak (Q), non-mitochondrial oxygen consumption (R), and spare respiratory capacity (S). (n = 4-6 biological replicates). Data are presented as mean ± SEM. Statistical significance was determined by unpaired two-tailed Student’s *t* test or one-way ANOVA. *p<0.05, **p<0.01, ***p<0.001, ****p<0.0001.

Single-nucleus multiomic profiling further corroborated these functional findings. Key genes involved in mitochondrial biogenesis and oxidative phosphorylation were upregulated in vhCOs (Figures S6D-S6G), including *PPARGC1A* (PGC-1α), a master regulator of mitochondrial metabolism known to drive cardiac maturation;^48,49^ Citrate synthase (*CS*), the rate-limiting enzyme of the tricarboxylic acid cycle (TCA) cycle, plays a pivotal role in mitochondrial energy production and is associated with cardiac maturation;^50^ and *NDUFA12*, a crucial accessory subunit of Complex I essential for ATP production.^51^ Furthermore, we found that vhCOs expressed much higher levels of key metabolic genes compared to hCOs, including those involved in fatty acid metabolism and amino acid metabolism (Figure S6G). Overall, these findings indicate that vascularization accelerates transcriptional, functional, and metabolic maturation of CMs.

### ECs adopt organotypic features in vhCOs

It has been indicated that the unique microenvironment within each organ critically drives the organotypic vascular endothelium identity.^52,53^ To evaluate the organ specificity of ECs in vhCOs, we first re-analyzed multiple independent human cell atlases^54,55^ to define organ-specific endothelial markers (Figure 3A). Notably, ECs from vhCOs acquired a cardiac-specific identity as indicated by the upregulated expression of cardiac-associated endothelial markers such as *LAMB1* and *MYADM* (Figure 3B). To further validate this finding, we isolated CDH5⁺ ECs from vhCOs and hBVOs and confirmed that vhCO-derived ECs showed significantly higher expression of *LAMB1* and *MYADM* compared with hBVO-derived ECs (Figures 3C and 3D). Consistently, whole-mount immunostaining further revealed more abundant Lamininβ1*^+^*CD31⁺ ECs in vhCOs than in hBVOs (Figure 3E), supporting the acquisition of a cardiac-specific endothelial phenotype.

**Figure 3.**
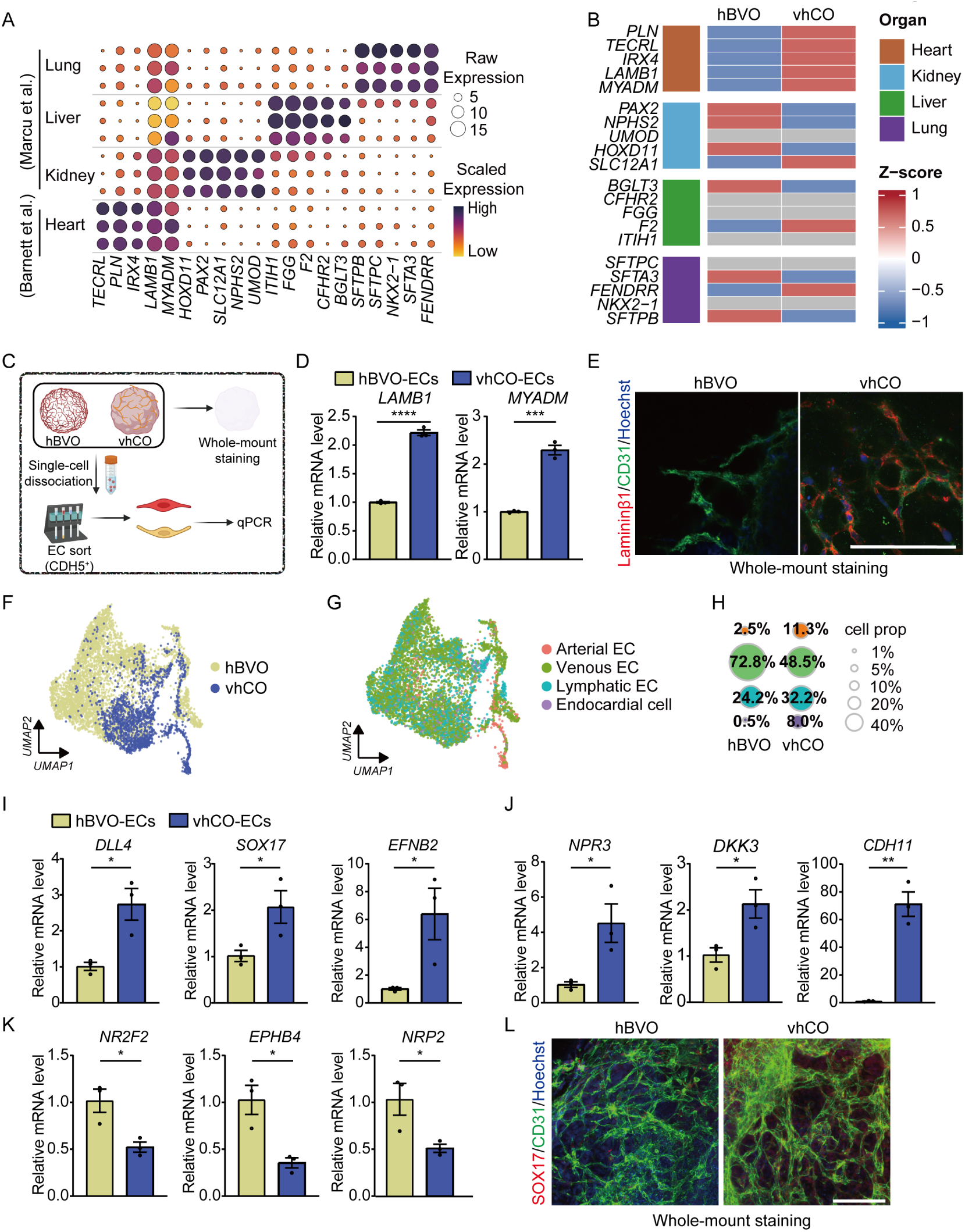
Vascularization establishes cardiac-specific endothelial cell identity in vhCOs. **(A)** Dot plot of organotypic EC markers generated based on scRNA-seq data of human fetal organs.^54,55^ Dot size denotes the raw expression level; color denotes scaled expression. **(B)** Heatmap of organ-specific EC markers in hBVO- and vhCO-derived ECs. **(C)** Schematic diagram of experimental workflow for whole-mount immunostaining and EC isolation. hBVOs and vhCOs were either processed for whole-mount immunostaining or dissociated, followed by CDH5⁺ EC sorting and qPCR analysis. **(D)** qPCR analysis of *LAMB1* and *MYADM* expression in sorted ECs, normalized to hBVO-derived ECs. (n = 3 biological replicates). **(E)** Representative whole-mount immunostaining of hBVOs and vhCOs for CD31 (green), Laminin β1 (red), and nuclei (blue). **(F)** UMAP visualization of integrated snRNA-seq data from hBVO- and vhCO-derived ECs. **(G)** Same UMAP as in (F), colored by endothelial subtypes (arterial, venous, lymphatic, and endocardial ECs). **(H)** Proportions of endothelial subtypes in hBVOs and vhCOs. Circle size reflects relative abundance. **(I-K)** qPCR analysis of arterial (I), endocardial (J), and venous EC markers (K) in sorted ECs, normalized to hBVO-derived ECs. (n = 3 biological replicates). **(L)** Representative whole-mount immunostaining of CD31 (green), SOX17 (red), and nuclei (blue). Scale bars, 50 μm. Data are presented as mean ± SEM. Statistical significance was determined by unpaired two-tailed Student’s *t* test. ***p<0.001, ****p<0.0001.

### ECs in vhCOs differentiate towards arterial and endocardial subtypes

Since snRNA-seq revealed that ECs in vhCOs and hBVOs clustered separately (Figure 3F), we next investigated whether the composition of EC subtypes differed between these organoids. Interestingly, the cardiac microenvironment led to a great enrichment of arterial ECs and endocardial cells in vhCOs, with approximately 5-fold and 10-fold increases in their respective proportion, accompanied by the reduced proportion of venous ECs (Figures 3G and 3H). These findings were further validated by qPCR analysis of isolated ECs from vhCOs and hBVOs. Expression of arterial markers (*DLL4*, *SOX17*, and *EFNB2*) and endocardial markers (*NPR3*, *DKK3*, and *CDH11*) were significantly upregulated, whereas venous markers (*NR2F2*, *EPHB4*, and *NRP2*) were significantly downregulated in vhCO-derived ECs (Figures 3I-3K). Immunofluorescence staining showed more SOX17^+^CD31^+^ ECs in vhCOs than in hBVOs (Figure 3L), consistent with the established role of *SOX17* in arterial specification.^56^ Collectively, these results indicate that cardiac microenvironment promotes the differentiation of hBVO-derived ECs toward arterial and endocardial lineages.

### *LAMA2* promotes arterial EC differentiation

To elucidate the intercellular signaling networks underlying the coordinated CM maturation and EC specification observed in vhCOs, we systematically analyzed the ligand-receptor interaction landscape among the seven major cell types identified in these organoids (Figure 4A). We first focused on signaling pathways regulating vascular formation and EC fate determination within vhCOs (Figure 4B). This analysis uncovered a dominant extracellular matrix (ECM)–integrin signaling module, in which collagens and Fibronectin (*FN1*) were dominantly secreted by FBs and myofibroblasts (MFBs), while laminins (e.g., *LAMA2*) was specific secreted by CMs, these ligands engage specific integrin receptors on ECs, including Integrin α1/β1 (*ITGA1/B1*), Integrin α2/β1 (*ITGA2/B1*), and Integrin α9/β1 (*ITGA9/B1*). This ECM-integrin signaling is known to drive vascular morphogenesis, governing EC sprouting, lumen formation, and vascular network assembly during embryonic development.^57–60^

**Figure 4.**
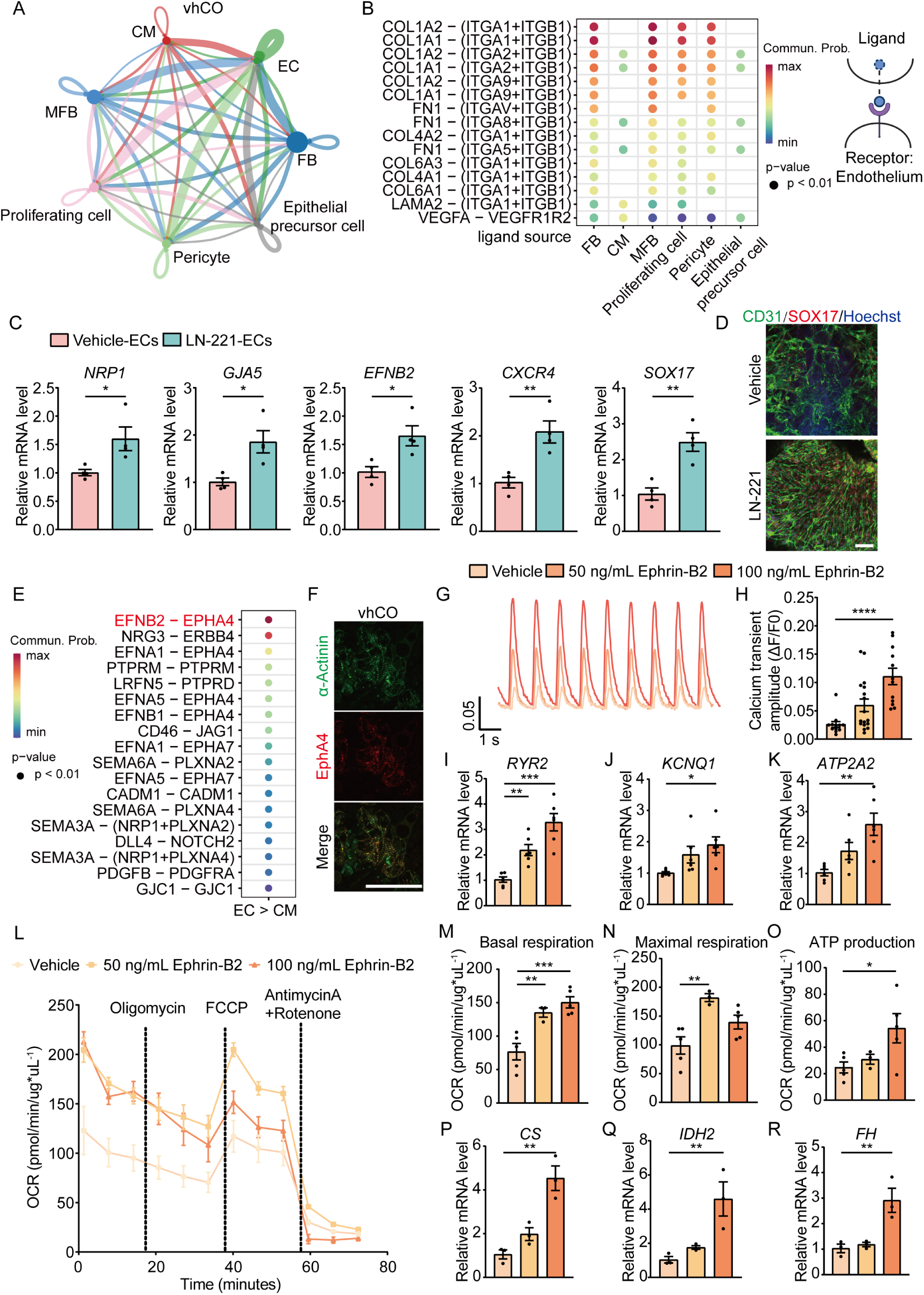
LN-221-mediated endothelial arterialization promotes cardiomyocyte maturation via EphrinB2 signaling. **(A)** Chord diagram showing global cell-cell communication networks within vhCOs. Line color and thickness indicate interaction pair and strength, respectively. **(B)** Dot plot of significant ligand-receptor interactions in which ECs act as signal receivers. **(C)** qPCR analysis of arterial EC marker genes in ECs isolated form hBVOs with or without Laminin-221 (LN-221) treatment, normalized to vehicle-treated ECs. (n = 4 biological replicates). **(D)** Representative whole-mount immunostaining of hBVOs for CD31 (green), SOX17 (red), and nuclei (blue). **(E)** Dot plot highlighting significant EC-to-CM ligand-receptor signaling pathways. **(F)** Representative vhCO section immunostaining for α-Actinin (green), EphA4 (red), and nuclei (blue). **(G)** Representative calcium transient traces of vhCOs treated with recombinant Ephrin-B2 (50, or 100 ng/mL). **(H)** Quantification of calcium transient amplitude. (n = 30-40 biological replicates). **(I–K)** qPCR analysis of *RYR2* (I), *KCNQ1* (J), and *ATP2A2* (K) expression in vhCOs following Ephrin-B2 treatment. (n = 6 biological replicates). **(L)** Seahorse extracellular flux analysis of OCR in vhCOs following Ephrin-B2 treatment. **(M-O)** Quantification of basal respiration (M), maximal respiration (N), and ATP production (O). (n = 4-6 biological replicates). **(P–R)** qPCR analysis of mitochondrial metabolic genes in vhCOs following Ephrin-B2 treatment. (n = 3 biological replicates). Scale bars, 50 μm. Data are presented as mean ± SEM. Statistical significance was determined by unpaired two-tailed Student’s *t* test or one-way ANOVA. *p<0.05, **p<0.01, ***p<0.001, ****p<0.0001.

Notably, as Collagen I and Collagen IV are exogenous components of the embedding matrix, while Fibronectin and Collagen VI are secreted by FBs in both hBVOs and vhCOs, they are unlikely to account for the endothelial phenotypic switching observed in vhCOs. We therefore focused on *LAMA2* in vhCO CMs, encoding the Laminin α2 chain of Laminin-221 (LN-221), a key basement membrane component of cardiac tissue (Figure 4B).^61^ LN-221 has been shown to promote differentiation of ESCs toward CM lineage and to improve cardiac function following cell transplantation in ischemic models,^62,63^ however, its role in endothelial specification has not been defined. To directly study this, we cultured hBVOs in the presence of LN-221 and observed a robust induction of arterial endothelial identity compared to controls. This was evidenced by significant transcriptional upregulation of canonical arterial EC markers including *NRP1*, *GJA5*, *CXCR4*, *EFNB2*, and *SOX17* (arterial-specifying transcription factor) (Figure 4C). Immunofluorescence staining further confirmed that LN-221 treatment increased the number of SOX17^+^CD31^+^ ECs in hBVOs (Figure 4D). Together, these data demonstrate that LN-221 drives ECs toward arterial cell fate in vhCOs.

### Arterial EC niche in vhCOs promotes CM maturation via Ephrin-B2-EphA4 signaling pathway

To investigate how EC-derived signals contribute to CM maturation, we next examined ligand-receptor interactions targeting CMs in vhCOs (Figure 4E). Among these, *EFNB2*—a gene encoding canonical arterial signaling ligand expressed by ECs—was significantly upregulated in vhCOs (Figure 3I) or following LN-221 treatment (Figure 4C), leading us to hypothesize that arterialized ECs establish a specialized paracrine niche that promotes CM maturation. This predicted interaction was spatially supported by the expression of EphA4—the receptor of Ephrin-B2—in CMs within vhCOs (Figure 4F). These observations suggest that LN-221-induced endothelial arterialization enables EC-CM communication via the Ephrin-B2-EphA4 axis, a pathway essential for cardiac lineage development.^64,65^

Consistent with this model, treatment of hCOs with recombinant Ephrin-B2 resulted in a dose-dependent enhancement of calcium handling, as reflected by increased calcium transient amplitudes compared to controls (Figures 4G and 4H). These functional enhancements were accompanied by elevated expression of calcium handling genes such as *RYR2* and *ATP2A2* and electrophysiological maturation related gene *KCNQ1* (Figures 4I–4K). Moreover, Seahorse metabolic analysis revealed that Ephrin-B2 treatment significantly increased basal and maximal mitochondrial respiration, as well as ATP production (Figures 4L–4O). This metabolic maturation was further supported by the upregulation of key TCA cycle genes including *CS*, *IDH2*, and *FH* (Figures 4P–4R). Together, these findings demonstrate that LN-221-mediated endothelial arterialization, establishes an *EFNB2-EPHA4* signaling axis that enhances both electrophysiological and metabolic maturation of CMs, thereby coupling vascular specialization with CM functional maturation in vhCOs.

### CFZ induces a dose-dependent cardiotoxicity in vhCOs

Having established physiologically relevant vhCOs, we next investigated CFZ-induced cardiovascular toxicity. To assess the specific contribution of vascularization to drug sensitivity, we compared vhCOs with hCOs and treated both models with two doses of CFZ. Notably, treatment with 500 nM CFZ resulted in a significant reduction in cell viability in vhCOs (Figure 5A), whereas no significant cytotoxicity was observed in hCOs (Figure S7A). This loss of viability was accompanied by a robust, dose-dependent induction of DNA damage, as indicated by increased level of phosphorylated H2A histone family member X (γ–H2AX) (Figure 5B). qPCR analysis revealed that CFZ markedly disrupted lineage-specific gene expression across multiple cardiac cell types, including CMs (*TNNT2* and *ACTA2*), ECs (*VWF*, *PECAM1*, *VEGFR1*, and *VEGFR2*), and pericytes (*PDGFRβ*) (Figure 5C). These changes indicate widespread cytotoxicity, loss of cellular identity, and coordinated multicellular apoptotic responses—phenotypes that are difficult to capture using 2D cultures or non-vascularized organoids. Immunofluorescence staining confirmed a dose-dependent increase in apoptosis in both α-Actinin⁺ CMs and CD31⁺ ECs (Figures 5D and 5E).

**Figure 5.**
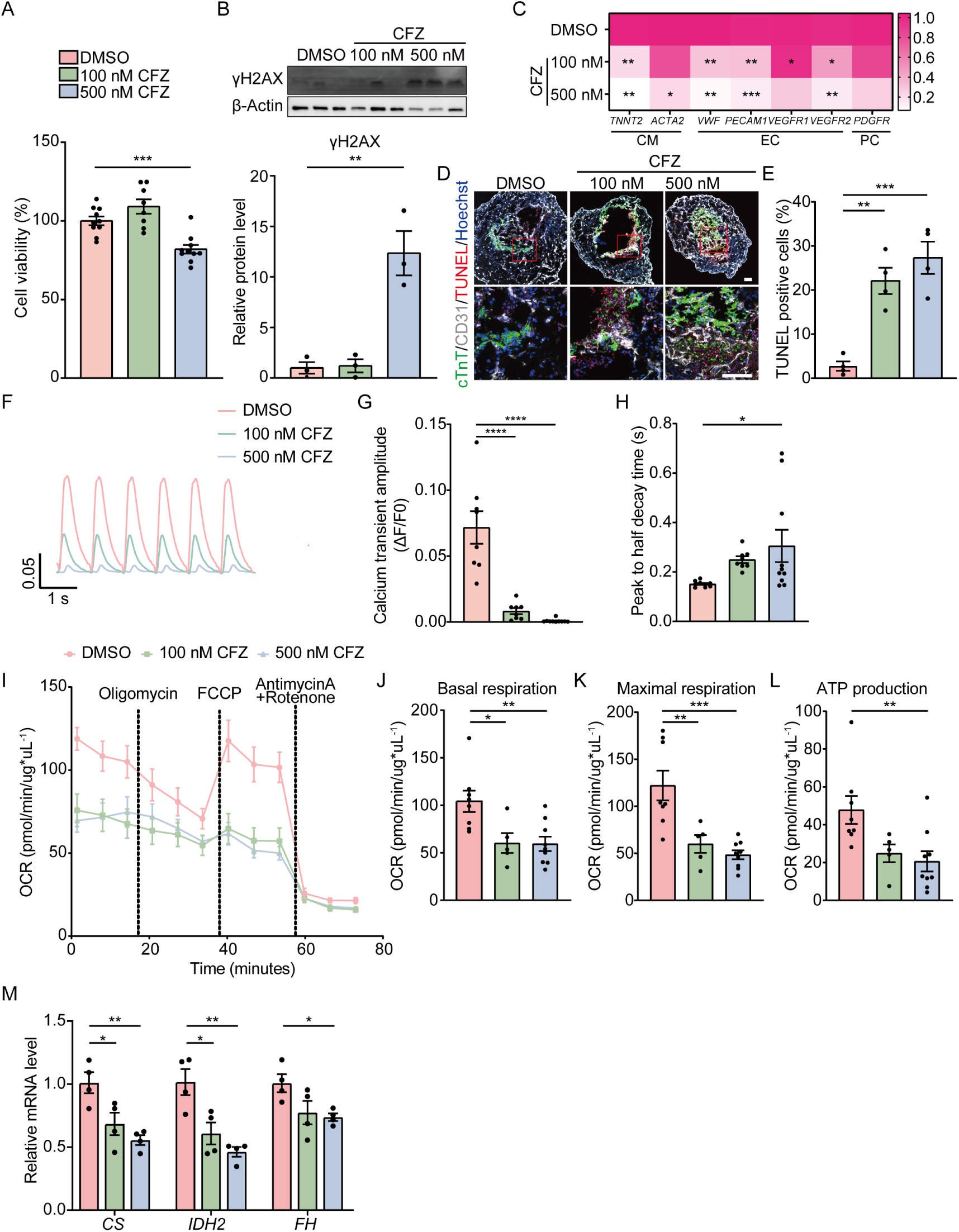
CFZ induces cardiotoxicity and mitochondrial dysfunction in vhCOs. **(A)** Cell viability of vhCOs treated with CFZ (100 or 500 nM) for 24 h. (n = 8-10 biological replicates). **(B)** Representative immunoblot (top) and quantification (bottom) of γH2AX in vhCOs. (n = 3 biological replicates). **(C)** Heatmap of relative expression of CM- and EC-specific marker genes following CFZ treatment, normalized to DMSO controls. (n = 3 biological replicates). **(D)** Representative fluorescence images and **(E)** quantification of TUNEL^+^ apoptotic cells in vhCO sections following CFZ treatment. (n = 4 biological replicates). **(F)** Representative calcium transient traces from vhCOs treated with CFZ. **(G-H)** Quantification of calcium transient amplitude (G) and peak to half decay time (H). (n = 8-10 biological replicates). **(I)** OCR of vhCOs treated with CFZ. **(J-L)** Quantification of basal respiration (J), maximal respiration (K), and ATP production (L). (n = 5-9 biological replicates). **(M)** qPCR analysis of mitochondrial metabolic genes (*CS*, *IDH2*, and *FH*) in vhCOs treated with CFZ. (n = 3 biological replicates). Scale bars, 100 μm. Data are presented as mean ± SEM. Statistical significance was determined by unpaired two-tailed Student’s *t* test or one-way ANOVA. *p<0.05, **p<0.01, ***p<0.001.

Given the central role of calcium handling in cardiac function, we next examined the effect of CFZ on calcium dynamics. CFZ exposure caused aberrant calcium transients in vhCOs, characterized by significantly reduced calcium transient amplitudes (Figures 5F and 5G, Supplementary Video S2) and prolonged calcium decay time (Figure 5H). In contrast, these calcium-handling defects were not observed in hCOs (Figures S7B and S7C). These findings indicate that vhCOs uniquely capture CFZ-induced calcium handling dysregulation, underscoring the necessity of vascularization for faithfully modeling the arrhythmogenic phenotypes observed clinically.^66^

Given that mitochondrial dysfunction is a known driver of calcium dysregulation,^67,68^ we next examined whether CFZ also induced mitochondrial dysfunction in vhCOs. Indeed, Seahorse metabolic analysis demonstrated significant reductions in basal respiration, maximal respiration, and ATP production following CFZ exposure (Figures 5I-5L). This metabolic dysfunction was further evidenced by the dose-dependent downregulation of critical TCA cycle genes (*CS*, *IDH2*, and *FH*) (Figure 5M), consistent with the observed mitochondrial dysfunction in animal models and MM patients.^69–71^ In contrast, hCOs exhibited no significant changes in mitochondrial respiration or ATP production following CFZ treatment (Figures S7D–S7G).

Taken together, these results demonstrate vhCOs as a robust and physiologically relevant *in vitro* model for dissecting drug-induced cardiotoxicity. Our data indicate that CFZ induces cardiotoxicity through multiple convergent mechanisms, including DNA damage, direct cytotoxicity, mitochondrial dysfunction, disruption of calcium homeostasis—pathological changes that are incompletely captured by conventional *in vitro* cardiac models.

### CFZ induces ATF4-dependent ER stress and IL8 expression in vhCOs

To further elucidate the molecular mechanisms underlying CFZ-associated cardiotoxicity, we conducted single-nucleus multiomic profiling on vhCOs treated with 500 nM CFZ. This analysis revealed a pronounced shift in FB heterogeneity: the homeostatic FB1 population decreased from 42.1% to 28.8%, whereas a distinct “stress-responsive” FB2 population expanded dramatically from 0.3% to 46.8% (Figures 6A and 6B). Consistent with this phenotypic transition, RNA velocity analysis delineated a clear trajectory from FB1 toward FB2 following CFZ exposure (Figure 6C).

**Figure 6.**
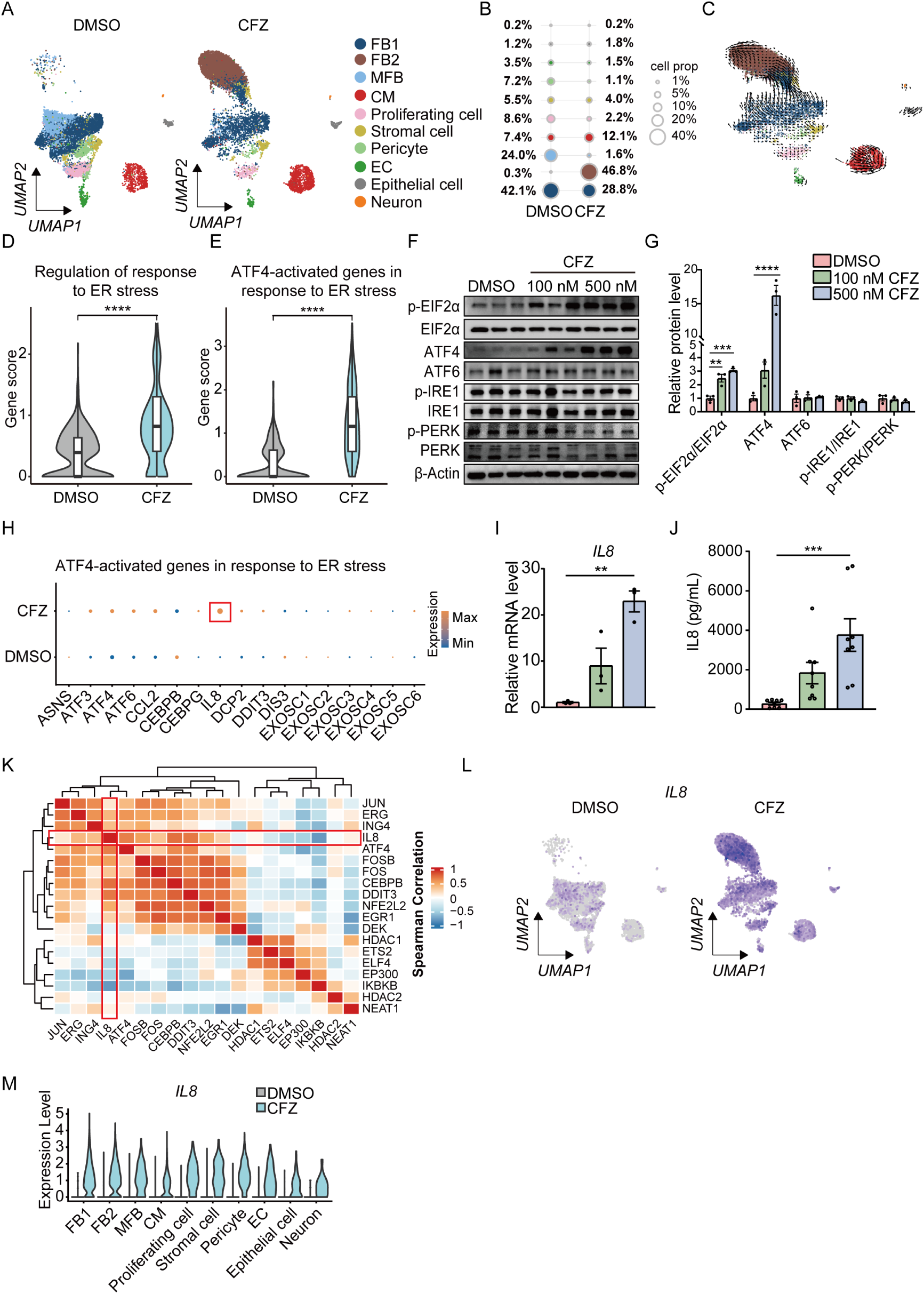
CFZ activates ER stress–responsive ATF4 signaling and induces IL8 expression in vhCOs. **(A)** UMAP visualization of snRNA-seq data from DMSO- and CFZ-treated vhCOs, colored by annotated cell type. **(B)** Relative cellular composition of vhCOs following CFZ treatment. **(C)** RNA velocity analysis projected onto the UMAP of vhCOs treated with CFZ. **(D–E)** Violin plots showing single-nucleus gene-set enrichment scores for “Regulation of response to ER stress” (D) and “ATF4-activated genes in response to ER stress” (E). **(F)** Representative immunoblots of ER stress pathway components in vhCOs treated with CFZ (100 or 500 nM). **(G)** Quantification of immunoblot data shown in (F). (n = 3 biological replicates). **(H)** Dot plot of representative ATF4 targeting genes. Dot size indicates the fraction of expressing cells; color denotes scaled average expression. **(I)** Relative IL8 mRNA expression measured by qPCR after CFZ treatment, normalized to DMSO. (n = 3 biological replicates). **(J)** Secreted IL8 protein levels in vhCO conditioned media measured by ELISA. (n = 8 biological replicates). **(K)** Heatmap of pairwise Spearman correlations between *ATF4*, *IL8*, and selected transcriptional regulators at the single-cell level. **(L)** UMAP visualization of IL8 transcript abundance in vhCOs treated with CFZ. **(M)** Violin plots of IL8 expression across major cell types. Data are presented as mean ± SEM. Statistical significance was determined by one-way ANOVA. **p<0.01, ***p<0.001, ****p<0.0001.

To identify the molecular drivers underlying this shift, we next examined stress-response signaling pathways. Gene set scoring showed a robust activation of the unfolded protein response (UPR), with significantly elevated enrichment scores for “Regulation of response to ER stress” and, more specifically, “ATF4-activated genes” in CFZ-treated vhCOs (Figures 6D and 6E). To validate these transcriptional signatures at the protein level, we examined three canonical UPR pathways. Western blot analysis revealed a dose-dependent increase in phosphorylated eIF2α (p-eIF2α) relative to total eIF2α and a concomitant upregulation of the downstream effector ATF4 (Figures 6F and 6G). These results indicate that the eIF2α-ATF4 axis represents the dominant ER stress pathway activated by CFZ.

We next examined how this ER stress response translated into downstream pathological signaling. Analysis of ATF4 transcriptional targets revealed a robust induction of ER stress-responsive genes following CFZ exposure, including canonical markers such as *DDIT3*, *ASNS*, and *CEBPB* (Figure 6H). Notably, *DDIT3* is a key pro-apoptotic mediator that is frequently co-activated with *ATF4* during ER stress.^72,73^ The upregulation of both *ATF4* and *DDIT3* following CFZ treatment (Figures 6F and 6G, and Figure S7H) indicates that CFZ-induced apoptosis is driven by ER stress pathway. Interestingly, within the ATF4-responsive gene set, we observed a selective upregulation of pro-inflammatory mediators, most prominently *CXCL8* (IL8) and C-C Motif Chemokine Ligand 2 (CCL2) (Figure 6H). Subsequent validation by qPCR and ELISA confirmed a significant, dose-dependent increase in IL8 expression and secretion in CFZ-treated vhCOs (Figures 6I and 6J). However, no significant changes in secreted IL8 were observed in either hCOs or hBVOs (Figures S7D and S7E), indicating a specific pro-inflammatory response in vhCOs. Furthermore, no significant changes in CCL2 expression or secretion were detected in vhCOs following CFZ exposure (Figures S7I and S7J). To further link IL8 induction to ER stress signaling, we performed Spearman correlation analysis across the transcriptomic dataset (Figure 6K). IL8 expression displayed a strong positive correlation with *ATF4* and other ER stress-response transcription factors. UMAP visualization and cell-type-resolved analysis further showed that IL8 expression increased across multiple cardiac cell types after CFZ treatment, but was most pronounced in FB1, FB2, and MFB populations (Figures 6L and 6M), highlighting FB-lineage specific sensitivity to inflammatory stress. These findings indicate that CFZ-induced inflammatory signaling depends on synergistic crosstalk among multiple cardiac lineages, underscoring the unique capacity of vhCOs to model stress amplification.

Together, these data demonstrate that CFZ triggers a robust ER stress response in vhCOs, characterized by activation of the eIF2α-ATF4 signaling, which in turn drives IL8-mediated inflammatory response across multiple cardiac cell populations.

### 4-PBA attenuates CFZ-induced ER stress and vascular disruption

To determine whether pharmacological suppression of ER stress could rescue CFZ-induced pathology, vhCOs were pretreated with the chemical chaperone and ER stress inhibitor 4-PBA prior to CFZ exposure. 4-PBA exhibited no detectable cytotoxicity and significantly prevented CFZ-induced cell death (Figure 7A). Consistent with this effect, CFZ-induced phosphorylation of eIF2α and upregulation of ATF4 were significantly suppressed by 4-PBA (Figures 7B-7E). We next investigated whether 4-PBA could alleviate CFZ-induced vascular toxicity. As expected, CFZ treatment caused substantial vascular disruption, as visualized by CD31 immunostaining (Figure 7F). Quantitative analysis revealed that 4-PBA pretreatment substantially rescued the CFZ-induced vascular injury, as indicated by increases in multiple vascular parameters, including total vessel area (Figure 7G), vessel area percentage (Figure 7H), total vessels length (Figure 7I), and total number of junctions (Figure 7J).

**Figure 7.**
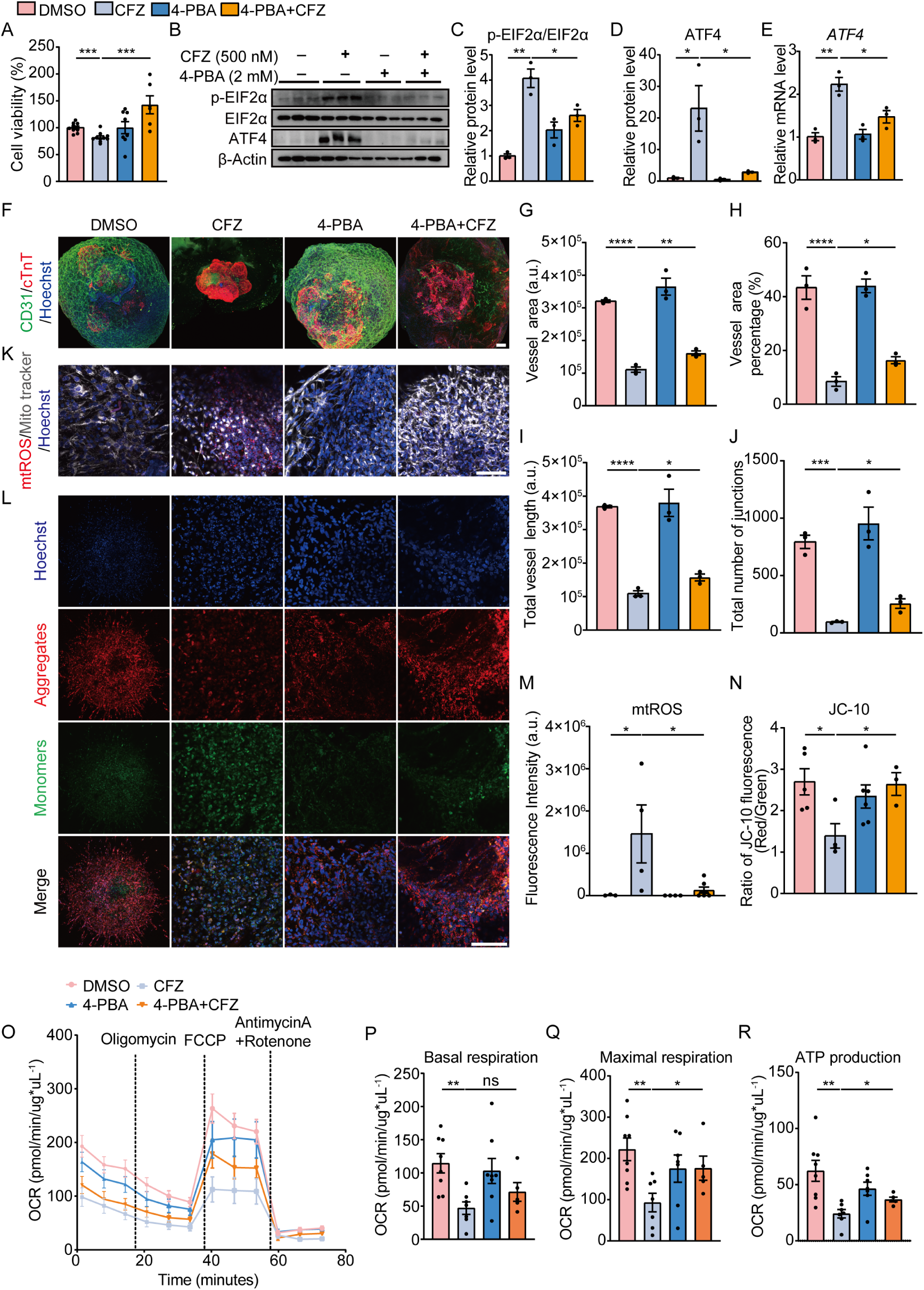
4-PBA attenuates CFZ-induced cardiotoxicity by suppressing ER stress and restoring mitochondrial function. **(A)** Cell viability of vhCOs treated with DMSO, CFZ (500 nM), 4-PBA (2 mM), or CFZ + 4-PBA for 24 h. (n = 6 biological replicates). **(B)** Representative immunoblots of p-eIF2α, total eIF2α, ATF4, and β-actin. **(C-D)** Quantification of p-eIF2α/eIF2α ratio (C) and ATF4 protein levels (D). (n = 3 biological replicates). **(E)** qPCR analysis of ATF4 mRNA expression under indicated conditions. (n = 3 biological replicates). **(F)** Representative immunofluorescence images of vhCOs stained for CD31 (green), cTnT (red), and nuclei (blue). **(G-J)** Quantification of vascular network parameters, including total vessel area (G), vessel area percentage (H), total vessel length (I), total number of junctions (J) analyzed from CD31-stained images using AngioTool. (n = 3 biological replicates). **(K)** Representative fluorescence images of mitochondria (MitoTracker), mitochondrial ROS (mtROS), and nuclei. **(L)** Representative JC-10 staining showing mitochondrial membrane potential. **(M-N)** Quantification of mtROS level (M) and JC-10 red/green fluorescence ratio (N). (n = 4-5 biological replicates). **(O)** OCR of vhCOs under indicated treatment. (n = 5-8 biological replicates). **(P-R)** Quantification of basal respiration (P), maximal respiration (Q), and ATP production (R). Scale bars, 100 μm. Data are presented as mean ± SEM. Statistical significance was determined by one-way ANOVA. *p<0.05, **p<0.01, ***p<0.001, ****p<0.0001.

Given the established interplay between ER stress and mitochondrial dysfunction, we further investigated the effect of 4-PBA on CFZ-induced mitochondrial dysfunction. mtROS staining revealed a pronounced CFZ-induced increase in mtROS, which was significantly attenuated by 4-PBA treatment, accompanied by restoration of mitochondrial membrane potential (Figures 7K-N). Consistent with these findings, Seahorse mitochondrial stress testing demonstrated that 4-PBA treatment could improve mitochondrial function, as evidenced by increased mitochondrial basal respiration, maximal respiration, and ATP production (Figures 7O-7R).

Collectively, these findings indicate that 4-PBA effectively mitigates CFZ-induced cardiotoxicity and vascular injury by suppressing ER stress and preserving mitochondrial function, thereby providing robust protection against CFZ-induced myocardial and vascular dysfunction in vhCOs.

## DISCUSSION

In this study, we reported a robust and faithful approach to generate vhCOs by assembling hCOs with hBVOs, resulting in branched vascular networks that were deeply integrated within cardiac tissues. Single-nucleus multiomic profiling revealed that the cellular composition of vhCOs closely mirrored that of the human fetal heart at 17 PCWs, encompassing CMs, ECs, FBs, and mural cell populations with appropriate lineage identities. This integrated multicellular niche enabled reciprocal cell-cell communications that drove both CM maturation and EC specification, yielding organoids that more faithfully recapitulated human cardiac development than conventional non-vascularized models.

Unlike previous vascularization strategies that rely on addition of ECs in cellular mixtures or geometrically confined co-development,^19,20^ assembled vhCOs supported extensive self-organization, allowing vascular plexuses to penetrate the myocardial core. This effectively reduced central apoptosis and sustained long-term culture. Importantly, incorporation of cardiac-specific vasculature emerged as an active driver of CM maturation. We observed a time-dependent progression of CM transcriptional, functional, and metabolic maturation following assembly, leading to CM state that closely resembled those of the human fetal heart at 17 PCWs, substantially more advanced than previously reported.^20^ These findings extend previous observations that EC–CM interactions promote CM maturation and demonstrate that vascular integration is required to unlock this developmental trajectory *in vitro*.^19^

A central challenge in the field has been the generation of organotypic ECs that faithfully reflect tissue-specific identity and function. Consistent with the role of local microenvironmental cues in EC specification,^54,74–77^ prolonged integration of hBVOs within a cardiac microenvironment promoted acquisition of cardiac-specific EC identities in vhCOs. ECs were enriched for *bona fide* cardiac endothelial markers, including *LAMB1* and *MYADM*, and exhibited transcriptional features distinct from pan-endothelial states.^21^ Lineage trajectory analysis further revealed that arterial and endocardial ECs emerged from venous EC intermediates, recapitulating the venous-to-arterial EC transition observed during blood vessel development.^77–82^ These findings established endothelial arterialization as one of hallmarks of vascular maturation in vhCOs,^83^ and demonstrated that our approach generated physiologically relevant and organotypic cardiac vasculature.

To define the molecular basis of vascular-driven cardiac maturation, we integrated single-nucleus multiomic data with cell-cell communication analysis, revealing ECM and Ephrin signaling as key regulators of arterial EC and CM states. We identified Laminin-α2 encoded by *LAMA2*, a core subunit of LN-221 that is the most abundantly expressed laminin in the basement membrane of mature cardiac muscle,^61^ emerged as a central cue promoting arterial EC specification. LN-221 enhanced expression of arterial markers such as *DLL4*, *CXCR4*, *SOX17*, and *EFNB2*, highlighting matrix-mediated control of endothelial fate. Ephrin-B2, a canonical arterial identity gene with established roles in vascular development and angiogenesis, and cardiac looping,^84–87^ further functioned as a bidirectional signal linking endothelial arterialization to CM maturation as shown in our study. These findings uncover a previously underappreciated pro-maturation role of the arterial endothelial niche in human heart development.

The absence of vasculature in existing hCO models has fundamentally limited their capacity to model cardiotoxicity induced by drugs that often affect multiple cell types in the heart. Using our vhCO platform, we recapitulated the multifaceted cardiotoxicity of CFZ, which is clinically associated with severe cardiovascular adverse events.^28–35^ CFZ induced dose-dependent CM injury characterized by reduced cell viability, increased DNA damage, and impaired calcium handling. Crucially, CFZ also disrupted endothelial and pericyte identities, indicating widespread vascular injury. These multicellular phenotypes closely align with clinical observations and underscore the importance of vascular integration for cardiotoxicity modeling.

Mechanistic dissection revealed that CFZ-induced cardiotoxicity arises from a cascade of interconnected molecular events centered on mitochondrial dysfunction and ER stress. CFZ impaired mitochondrial integrity as evidenced by increased mtROS, membrane potential depolarization, and suppression of oxidative phosphorylation. Further analysis revealed that activation of the eIF2α-ATF4 axis drove the FB state transition from the homeostatic FB1 to the stress-responsive phenotype (FB2) and robust secretion of the pro-inflammatory cytokine IL8. Importantly, this inflammatory response was largely blunted in non-vascularized hCOs or hBVOs, indicating that vascularized cardiac multicellular microenvironment is required to reveal this pathogenesis. Following these findings, we identified the FDA-approved ER stress inhibitor 4-PBA as an effective intervention that attenuated ATF4-mediated stress response, preserved mitochondrial bioenergetics, and maintained vascular integrity in vhCOs. These findings corroborate previous reports highlighting the potential of 4-PBA to restore mitochondrial function in the cardiac aging model^88^ and sepsis-induced cardiomyopathy.^89^

In conclusion, this study establishes vhCOs as a physiologically faithful platform for modeling cardiac development and disease. By coaxing organotypic vasculature in cardiac microenvironment, vhCOs enable study of endothelial-cardiomyocyte crosstalk, undercover vascular-dependent disease mechanisms, and provide a robust platform for therapeutic testing.

### Limitations of the study

While vhCOs exhibit an advanced maturation state relative to existing hCO models, they retain characteristics akin to the human fetal heart and do not yet achieve full adult-like structural and functional maturity. This limitation may, in part, reflect the absence of specific lineage populations, including immune cells (e.g., macrophages) and autonomic neurons, which are known to play a critical role in cardiac homeostasis, remodeling, and disease progression *in vivo*. Recent studies have shown that macrophages promote organoid maturation and functionality.^90–92^ The lack of these populations limits the capacity of vhCOs to fully model intercellular interactions and disease mechanisms. Future studies incorporating additional cardiac-relevant lineages will be necessary to further enhance cellular complexity and physiological relevance. Despite the presence of an organized and extensive vascular network, vhCOs are generated in the absence of active hemodynamic flow. The lack of perfusion restricts exposure to biomechanical cues, such as shear stress, which are known to affect cardiac maturation and function. Incorporation of controllable microfluidic perfusion systems has been demonstrated to improve organoid maturation and function and may address this limitation.^93–97^ Finally, the limited availability of MM patients’ samples limited the mechanistic validation in this study. Future investigations using human primary tissues will be important to further substantiate these findings and to improve the translational feasibility of vhCOs for disease modeling and drug discovery.

## RESOURCE AVAILABILITY

### Lead contact

Further information and requests for resources and reagents should be directed to and will be fulfilled by the lead contact, Joe Z Zhang.

### Materials availability

This study did not generate new unique reagents.

### Data and code availability

The data and code are available from the corresponding author on reasonable request. Any additional information required to reanalyze the data reported in this paper is available from the lead contact upon request.

## ACKNOWLEDGEMENTS

This study was supported by the National Natural Science Foundation of China (W2432052 and 82370311) and grants from the Science and Technology Development Fund (FDCT) of Macau (FDCT/0004/2021/AKP and FDCT/0038/2020/AFJ) and the University of Macau internal grant (MYRG-GRG2024-00273-FHS and SRG2019-00177-FHS). Some illustrations were created with BioRender.com. hiPSC cell lines used in this study were kindly provided by Stanford Cardiovascular Biobank. We appreciate the support of SZBL Biomedical Research Core Facilities and Laboratory Animal Center. We thank Dr. Kai Yang from the Bioimaging Core of SZBL for technical assistance with image processing and 3D reconstruction.

## AUTHOR CONTRIBUTIONS

Y.Y. and L.Z. contributed equally to this work. J.Z.Z., N.-Y.S., Y.Y., and L.Z. conceived and designed the project. Y.Y., H.T., Y.Z., H.X., and H.W. conducted the experiments. L.Z. conducted bioinformatical analysis. All authors analyzed and discussed the data. Y.Y., L.Z., N.M., N.-Y.S., and J.Z.Z wrote the manuscript.

