## Supplementary information for "Vascularized Human Cardiac Organoids Reveal Endothelial–Cardiomyocyte Crosstalk and Mechanisms of Carfilzomib-Induced Cardiotoxicity"

**Supplementary Figure Legends**

**Figure S1. Generation of hCOs. (A)** Schematic diagram of the differentiation protocol used to generate hCOs from hiPSCs. EBs: embryonic bodies. **(B)** Percentage of beating organoids derived from four individual hiPSC lines across four independent differentiation batches. **(C)** Temporal expression of key cardiac marker genes during hCO differentiation. (n = 3 biological replicates). **(D)** Representative whole-mount immunostaining of D12 hCOs showing CM markers (α-Actinin, CX43, and cTnT) and the endothelial marker (CD31). Scale bars, 100 µm. Data are presented as mean ± SEM. Statistical significance was determined by one-way ANOVA. *p<0.05, ***p<0.001, ****p<0.0001.

**Figure S2. Single-nucleus multiomic profiling reveals cellular diversity in hCOs. (A)** UMAP visualization of integrated snRNA-seq data from D15 and D42 hCOs, identifying 16 transcriptionally distinct clusters. Major cell types are annotated. Inset shows D15 (pink) and D42 (purple) cells. **(B)** Cellular composition of hCOs. **(C)** UMAP feature plots showing expression of representative marker genes in CMs, CPCs, and cardiac neural crest cells (CNC), and epithelial precursor cells. Color scale indicates normalized expression. **(D)** Representative genomic region showing snATAC-seq chromatin accessibility tracks for marker genes in indicated cell types in (C).

**Figure S3. Generation and functional characterization of hBVOs. (A)** Schematic diagram of the differentiation protocol used to generate hBVOs from hiPSCs. **(B)** Temporal expression of key endothelial and mural cell marker genes during hBVO differentiation. (n = 3 biological replicates). **(C)** Representative whole-mount immunostaining of D12 hBVOs showing the endothelial marker CD31 and mural cell markers PDGERβ and vimentin. Yellow box indicates higher-magnification view. **(D)** Schematic diagram of the experimental workflow for functional characterization of hBVOs, including transplantation under the kidney capsule of NSG mice to examine vascularization and dissociation of ECs for LDL uptake assays. **(E)** Representative fluorescence images of grafts showing mouse CD31 (mCD31, red) and human CD31 (hCD31, green) expression (left). Vascular anastomosis was assessed by intravenous injection of Texas Red–dextran and co-staining with hCD31 (right). **(F)** DiI-Ac-LDL uptake (red) by ECs *in vitro*. Scale bars, 100 μm. Data are presented as mean ± SEM. Statistical significance was determined by one-way ANOVA. *$p<0.05$, **$p<0.01$, ***$p<0.001$, ****$p<0.0001$.

**Figure S4. Single-nucleus multiomic profiling reveals cellular diversity in hBVOs. (A)** UMAP visualization of integrated snRNA-seq data from hBVOs, identifying five transcriptionally distinct clusters. Major cell types are annotated. **(B)** Cellular composition of hBVOs. **(C)** UMAP feature plots showing expression of representative marker genes in five clusters. Color scale indicates normalized expression. **(D)** Representative genomic region showing snATAC-seq chromatin accessibility tracks for marker genes in indicated cell types in (C).

**Figure S5. Optimization of vascularization strategies and culture conditions for vhCOs. (A)** Representative whole-mount immunostaining of vhCOs for cTnT (red) and CD31 (green) in three different medium conditions: CDM, RPMI, and a 1:1 RPMI+CDM mixture. Yellow boxes indicate higher-magnification views. **(B)** Representative whole-mount immunostaining of vhCOs generated using two fusion ratios: hBVO to hCO at 1:1 (top) and 1:2 (bottom). **(C–E)** Quantitative analysis of vascular network formation in vhCOs, including total vessel length (C), vessel area (D), and total number of junctions (E) (n = 3-5 biological replicates). **(F-G)** Three-dimensional surface rendering shows CMs (cTnT, red) and ECs (CD31, green) (F) and the lumen formation of ECs (G) in vhCO. Scale bars, 100 μm. Data are presented as mean ± SEM. Statistical significance was determined by one-way ANOVA. *p<0.05, **p<0.01.

**Figure S6. Single-nucleus multiomic profiling reveals enhanced electrophysiological and metabolic maturation in vhCOs. (A–F)** Violin plots (top) and chromatin accessibility tracks (bottom) showing gene expression and open chromatin at loci of indicated genes in hCOs and vhCOs at D15 and D42. **(G)** Heatmap of relative expression of key metabolic genes related to amino acid metabolism, fatty acid metabolism, mitochondrial dynamics, and TCA cycle & ETC in vhCOs compared with hCOs.

**Figure S7. CFZ-induced cardiotoxicity is negligible in hCOs. (A–G)** Minimal cardiotoxicity in hCOs treated with CFZ (100 or 500 nM). Quantification of cell viability (A) (n = 7-8 biological replicates), calcium transient amplitude (B) (n = 7-12 biological replicates), and peak-to-half decay time (C) (n = 9-12 biological replicates), OCR of hCOs (D), quantification of basal respiration (E), maximal respiration (F), and ATP production (G). (n = 5-6 biological replicates). **(H-I)** qPCR analysis of *DDIT3* (H) and *CCL2* (I) expression in vhCOs (n = 3 biological replicates). **(J)** Secreted CCL2 protein levels in vhCO conditioned media measured by ELISA. (n = 3 biological replicates). **(K-L)** Secreted IL8 protein levels in hCO (K) and hBVO (L) conditioned media measured by ELISA. Data are presented as mean ± SEM. Statistical significance was determined by one-way ANOVA. ***$p<0.001$.

**Supplementary Tables**

**Table S1.** Sequence of primers used for qPCR.

**Table S2.** List of antibodies.

**Supplementary Videos**

**Video S1. 3D reconstruction of vascular networks of D42 vhCO.** This video presents a 3D volumetric rendering of a D42 vhCO, highlighting the branched and lumenized vascular network stained by CD31 (green) integrated with myocardial tissue stained by cTnT (red).

**Video S2. CFZ-induced calcium handling abnormalities in vhCOs.** Representative real-time calcium imaging recording of vhCOs following CFZ treatment.

**Table S1.** Sequence of primers used for qPCR.

| Gene | Forward primer | Reverse primer |
|---|---|---|
| *18S* | CGGCTACCACATCCAAGGAA | CCTGTATTGTTATTTTTCGTCACTACCT |
| *SM22* | TCCAGGTCTGGCTGAAGAATGG | CTGCTCCATCTGCTTGAAGACC |
| *PECAM1* | AGACGTGCAGTACACGGAAG | TTTCCACGGCATCAGGGAC |
| *VEGFR1* | AACGTGGTTAACCTGCTGGG | AGTGCTGCATCCTTGTTGAGA |
| *VEGFR2* | GAGGGGAACTGAAGACAGGC | GGCCAAGAGGCTTACCTAGC |
| *MYH7* | TCACCAACAACCCCTACGATT | CTCCTCAGCGTCATCAATGGA |
| *SOX2* | GCTACAGCATGATGCAGGACCA | TCTGCGAGCTGGTCATGGAGTT |
| *TBXT* | CCTTCAGCAAAGTCAAGCTCACC | TGAACTGGGTCTCAGGGAAGCA |
| *NKX2.5* | AAGTGTGCGTCTGCCTTTCCCG | TTGTCCGCCTCTGTCTTCTCCA |
| *GATA4* | TGCCGTTCATCTTGTGGTAG | CCGACACCCCAATCTCG |
| *TNNT2* | TTCACCAAAGATCTGCTCCTCGCT | TTATTACTGGTGTGGAGTGGGTGT |
| *MYH6* | GGAAGACAAGGTCAACAGCCTG | TCCAGTTTCCGCTTTGCTCGCT |
| *MYBPC3* | AACCTGTCAGCCAAGCTCCACT | CCACAATGGTGTCTGGTATGCG |
| *ACTA2* | CTATGCCTCTGGACGCACAACT | CAGATCCAGACGCATGATGGCA |
| *CNN1* | CCAACGACCTGTTTGAGAACACC | ATTTCCGCTCCTGCTTCTCTGC |
| *RYR2* | CTTGAGGTTGGCTTTCTGCCAG | TGTGCCAGCAAAGAGAGGAGCA |
| *KCNQ1* | CTCCGTGGTCTTCATCCAC | TCCTTCTCAGCCAGGTACACA |
| *MYADM* | CCAGTTCGATGAGAAGTATGGCG | CAGCCACATACGCCAGTAGGTT |
| *LAMB1* | GAGGTGTCTCAAGTGCCTGTAC | ACTGGCAGTCAGAGCCGTTACA |
| *DLL4* | CCAACTGTGGCAAACAGCAA | GCTCTTGTCACTGTGGGGAA |
| *SOX17* | TCTGCCTCCTCCACGAAG | ACGCCGAGTTGAGCAAGA |
| *EFNB2* | CTCCTCAACTGTGCCAAACCA | GGTTATCCAGGCCCTCCAAA |
| *NR2F2* | CGGGTGGTCGCCTTTATGG | ACAGGCATCTGAGGTGAACAG |
| *EPHB4* | CCACCGGGAAGGTGAATGTC | CTGGGCGCACTTTTTGTAGAA |
| *NRP2* | GCTGGCTATATCACCTCTCCC | TCTCGATTTCAAAGTGAGGGTTG |
| *NRP1* | GGCGCTTTTCGCAACGATAAA | TCGCATTTTTCACTTGGGTGAT |
| *CDH11* | GATCGTCACACTGACCTCGACA | CTTTGGCTTCCTGATGCCGATTG |
| *NPR3* | CAGTGGAGACTACGCCTTCTTC | TGACTGTCTGGAGGGACGAGTA |
| *DKK3* | GTGCATCATCGACGAGGACTGT | TGGTCTCCACAGCACTCACTGT |
| *GJA5* | AATCTTCCTGACCACCCTGCATGT | CAGCCACAGCCAGCATAAAGACAA |
| *CXCR4* | ACTACACCGAGGAAATGGGCT | CCCACAATGCCAGTTAAGAAGA |
| *ATP2A2* | GGACTTTGAAGGCGTGGATTGTG | CTCAGCAAGGACTGGTTTTCGG |
| *CS* | CACAGGGTATCAGCCGAACCAA | CCAATACCGCTGCCTTCTCTGT |
| *IDH2* | AGATGGCAGTGGTGTCAAGGAG | CTGGATGGCATACTGGAAGCAG |
| *FH* | CCGCTGAAGTAAACCAGGATTATG | ATCCAGTCTGCCATACCACGAG |
| *IL8* | GAGAGTGATTGAGAGTGGACCAC | CACAACCCTCTGCACCCAGTTT |

**Table S2.** List of antibodies.

| Antibody | Source | Catalog number |
|---|---|---|
| Mouse monoclonal anti PECAM1 | R&D | BBA7 |
| Rabbit polyclonal anti cTnT | Proteintech | 15513-1-AP |
| Purified anti-mouse CD31 | Biolegend | 102501 |
| Purified anti-human CD31 | Biolegend | 303102 |
| Mouse monoclonal anti PDGFRβ | Santa Cruz | sc-374573 |
| Rabbit monoclonal anti Vimentin | Abcam | ab92547 |
| Mouse monoclonal anti α-Actinin | Sigma | A7811 |
| Rabbit anti Connexin 43 | Affinity | AF0137 |
| Rabbit polyclonal anti LAMB1 | Proteintech | 23498-1-AP |
| Rabbit polyclonal anti EPHA4 | Proteintech | 21875-1-AP |
| Rabbit polyclonal anti SOX17 | Proteintech | 24903-1-AP |
| Rabbit monoclonal anti Phospho-eIF2α(Ser51) | Cell Signaling Technology | 3398 |
| Rabbit polyclonal anti eIF2α | Cell Signaling Technology | 9722 |
| Rabbit recombinant anti-ATF4 | Proteintech | 81798-2-RR |
| Rabbit polyclonal anti ATF6 | Proteintech | 24169-1-AP |
| Rabbit polyclonal anti Phospho-PERK (Thr982) | Affinity | DF7576 |
| Rabbit polyclonal anti PERK | Abcam | ab65142 |
| Rabbit anti Phospho-IRE1 (Ser724) | Affinity | AF7150 |
| Rabbit polyclonal anti IRE1 | Proteintech | 27528-1-AP |
| IRDye 680RD-conjugated polyclonal goat anti-Mouse IgG | LICORbio | 926-68070 |
| IRDye 800CW-conjugated polyclonal goat anti-Rabbit IgG | LICORbio | 926-32211 |
| Goat anti Rabbit IgG (H+L) Alexa Fluor 488 | Cell Signaling Technology | 4412 |
| Goat anti Rabbit IgG (H+L) Alexa Fluor 555 | Cell Signaling Technology | 4413 |
| Goat anti Mouse IgG (H+L) Alexa Fluor 488 | Cell Signaling Technology | 4408 |
| Goat anti Mouse IgG (H+L) Alexa Fluor 555 | Cell Signaling Technology | 4409 |
| Goat anti Mouse IgG (H+L) Alexa Fluor 647 | Thermo Fisher Scientific | A21236 |

### Methods

#### hiPSC maintenance

hiPSCs were routinely maintained in the E8 medium (Gibco, A1517001) on Matrigel-coated 6-well plates (Matrigel; Corning, 354230) as described previously (ref). Cells were dissociated with 0.5 mM EDTA (Thermo Fisher Scientific, 15575020) in PBS when reached 70–80% confluency. Dissociated cells were then passaged in the E8 medium supplemented with 10 μM Rho-associated protein kinase inhibitor Y-27632·2HCl (Selleck Chemicals Cat, S1049) and cultured for 24 h. All cell cultures were maintained within a humidified incubator at 37℃, 5% $CO_2$. All hiPSC-related work was approved by the Shenzhen Bay Laboratory (SZBL) Medical Ethics Committee (YL 2023-002 and YL 2025-017).

#### Animals

Immune-deficient NOD-SCID IL-2Rγnull (NSG) mice at the age of 8 weeks old were obtained from the Guangdong GemPharmatech Co.,Ltd for hBVO transplantation experiments. Both male and female mice were used in the study and we did not observe any gender effect on the results. All mice were housed under 12 h/12 h day and night cycle with water and food *ad libitum*. All animal procedures were approved by SZBL Animal Ethics Committee (D2022-072 and D2025-119).

#### Differentiation of hiPSCs into hBVOs

hBVOs were generated based on a protocol described by Wimmer et al.[20] with modifications. Briefly, hiPSCs were dissociated into single cells and seeded into anti-adherence rinsing solution (Stem Cell Technologies, 0-7010) -coated U-shaped bottom 96-well plates with 3000 cells/well. Following centrifugation for 3 min at 300× g at room temperature, aggregates are formed in the E8 medium supplemented with 10 μM Y-27632·2HCl. hBVOs differentiation was induced with a chemically defined medium (CDM). The CDM was prepared by mixing 245 ml of DMEM/F-12 (Gibco, 10565018), 245 ml of IMDM (Gibco, C12440500BT), 5 ml of chemically defined lipid concentrate (Gibco, 11905031), 5 ml of GlutaMAX (Gibco, 35050-061), 500 μl of Insulin-Transferrin-Selenium (Abmole, M19989), and 20 μl of α-monothioglycerol (Sigma-Aldrich, M6145). hiPSCs were induced for vascular lineage cell differentiation by continuous treatments with 12 μM CHIR99021 (LC Laboratories, C-6556) and 30 ng/mL BMP4 (PeproTech, 120-05ET) for 3 days. On day 3, medium was replaced by a cocktail of 100 ng/mL VEGF165 (PeproTech, 100-20), 100 ng/mL FGF2 (PeproTech, 100-18B), and 10 μM SB431542 (Selleck, 1067) in CDM for two days. hBVOs were then embedded into a 4:1 mixture of Collagen I-Matrigel solution. The Collagen I solution was adapted from Wimmer et al. [20] and recommendations by the manufacturers' protocol. To prepare 2 mL of 2.0 mg/mL collagen I solution, mix 300 μL of 0.1 N NaOH (Macklin, S817973), 125.2 μL of 5× DMEM (Vivacell, C3116-0500), 25.2 μL of HEPES (Beyotime, C0217), 20 μL of 7.5% sodium bicarbonate (GibcoCat, 25080094), 12.4 μL of Glutamax, 184 μL of Ham's F-12 (Gibco, 31765035) and 1.3 mL of 3 mg/ml PureCol Type I Collagen Solution (Advanced Biomart, 5005). The embedded organoids were then incubated with CDM medium supplemented with 100 ng/mL VEGF165 (PeproTech, 100-20) and 100 ng/mL FGF2 (PeproTech, 100-18B). The medium was replaced every three days throughout the differentiation period.

**Differentiation of hiPSCs into hCOs**

On day 0, hCOs were generated by seeding 5,000–10,000 cells/well into anti-adherence rinsing solution-coated U-shaped bottom 96-well plates in the E8 medium supplemented with 10 μM Y27632·2HCl. Aggregates were formed by centrifugation for 3 min at 300x g at room temperature. After two days, the medium was replaced with RPMI 1640 (Gibco, C11875500BT) supplemented with B27 minus insulin (Gibco, A18956-01) containing 10 ng/ml BMP4, 50 ng/ml activin A (PeproTech, 120-14P), and 6 μM CHIR-99021 for 48 h. On day 4, the medium was replaced with fresh differentiation medium. On day 5, the medium was changed to RPMI 1640 supplemented with 3 μM Wnt-C59 (Abmole, M3131) for two days. The medium was then replaced with differentiation medium for two days followed by changing to RPMI 1640 supplemented with B27 (Gibco, 17504044) for three days. The medium was changed every other day with maintenance medium until experiment.

**Generation of vhCOs**

To generate vhCOs, a day 7 hBVO and a day 12 hCO were collected, co-embedded in 20 μl Collagen I–Matrigel mixture as described above, and centrifuged for 3 min at 300x g at room temperature. The vhCO was then polymerized at 37 ℃ for 2 h. Following polymerization, a 1:1 mixture of CDM and RPMI 1640/B27 medium supplemented with 100 ng/ml VEGF165 and 100 ng/ml FGF2 was added to the vhCOs for maintenance.

**Single-cell suspension preparation**

Organoids were collected in a 50 mL falcon tube, washed with PBS, and then incubated in 0.5 mM EDTA/PBS buffer at 37℃ for 5 min. The digestion solution was subsequently replaced with PBS containing 2 mg/mL collagenase II (Worthington, LS004176), and organoids were incubated for 30 min at 37℃ on a shaker at 200 rpm. Following centrifugation at 300x g for 3 min at room temperature, cells were resuspended in RPMI 1640/B27 supplemented with 10% KnockOut serum replacement (KOSR, Thermo Fisher Scientific, 10828028). Cell suspensions with a cell density of 1,000 cells/μL and viability ≥90% were used for flow cytometry analysis or nuclear extraction prior to single-sell multiomic library preparation.

**Cell viability**

Cell viability was assessed using CellTiter-Glo® Luminescent Cell Viability Assay (Beyotime, C0061S), following the manufacturer's recommended protocols.

**Calcium transient recordings**

To record calcium transients in hCOs and vhCOs, organoids were incubated with Rhod-2 AM (2 μM, Thermo Fisher Scientific, R1245MP) in Tyrode's solution (Pricella, PB180340) for 30 min at 37℃. After loading, the solution was replaced with fresh Tyrode's solution and the organoids were incubated for an additional 30 min at 37℃ to allow for dye de-esterification. Imaging was performed at 44 frames per second on an inverted microscope (Nikon, Eclipse Ti2-E) equipped with a high-speed Andor Zyla sCMOS camera (Oxford Instruments) or IonOptix Optical Imaging System (IonOptix LLC). Calcium transients were recorded while organoids were electrically paced at 1 Hz and data were analyzed using customized MATLAB scripts.

**Quantitative real-time PCR (qPCR)**

Total RNA was extracted from organoids using the RNA extraction kit (Tiangen, DP451) and cDNA was prepared using a Takara Prime Script RT Reagent Kit (TaKaRa, RR037A) according to the manufacturers' instruction. qPCR was performed using Taq Pro Universal SYBR qPCR Master Mix (Vazyme Biotech, Q712-03) in a CFX384 Optical Reaction Module on C1000 Touch Thermal Cycler (Bio-Rad). The relative mRNA abundance of target genes was normalized to that of 18S mRNA obtained in the same sample. The primer sequences are shown in Table S1.

**Immunofluorescence staining of organoid cryosections**

Organoids were washed in PBS before fixing in 4% PFA for 1 h at room temperature. After washing three times in PBS, organoids were then embedded in OCT (Sakura, 4583) and snap frozen in an isopentane bath pre-cooled with liquid nitrogen and stored at −80℃. For immunostaining, samples were sectioned in slices of 8 μm thickness at −20℃ and were fixed with 4% PFA for 10 min, followed by permeabilized with 0.2% Triton X-100 for 30 min at room temperature. After blocking with 5% bovine serum albumin (BSA, Solarbio, A8020) for 1 h, the sections were incubated with primary antibodies overnight at 4℃. After washing three times with PBS, the sections were incubated with the secondary antibodies at a concentration of 1:400 at room temperature for 1 h protected from light. After staining with Hoechst 33342 (Thermo Fisher Scientific, 62249), samples were mounted with fluorescence mounting medium (Servicebio, G1401) and stored at 4℃ until imaging using Olympus FV3000 confocal microscope. The antibodies used for immunofluorescence staining are listed in Table S2

**Three-dimensional imaging of whole mount organoids**

After washing with PBS, the organoids were fixed with 4% PFA for 1 h at room temperature on a shaker at 200 rpm, followed by permeabilized with 0.2% Triton X-100 for 1 h. They were subsequently incubated with primary antibodies for 24 h, followed by incubation with secondary antibodies for 24 h, and then stained with Hoechst 33342 for 1 h at room temperature. Whole-organoid clearing was then performed using the iDISCO method as described online (https://idisco.info/) with some modifications. Briefly, organoids were dehydrated at room temperature by gradient methanol/$H_2O$ solutions (20%, 40%, 60%, 80%, and 100%, 1 h each) on a shaker at 200 rpm. To ensure complete removal of residual water, the organoids were washed in 100% methanol solution for 1 h. After dehydration, they were transferred to a glass bottle and incubated in 100% dichloromethane solution for 1 h at room temperature. Finally, organoids were incubated in 100% dibenzyl ether (Tokyo Chemical Industry, TCI-B0418) until they became optically transparent. Transparent organoids were then placed in a chamber filled with dibenzyl ether and imaged using an Olympus FV3000 microscope. Imaris software 9.0.1 (Oxford Instruments) was used to perform 3D reconstruction and 3D data analysis. The antibodies used for whole-mount staining are listed in Table S2.

**Quantification of the vascular network**

To quantify the vascular structure, organoids were immunostained with CD31 antibody and the images were analyzed using AngioTool software (https://github.com/imagejan/angiotool).

**Mouse kidney capsule transplantation**

D30 hBVOs were transplanted under the kidney capsule of 8-week-old immunodeficient NSG mice. In brief, mice were anesthetized, and the right kidney was exposed. After a small incision was made into the renal capsule, it was carefully lifted from the functional tissue to allow for insertion of the organoid. Once organoids were inserted, the wound was closed with electrocoagulation pen to avoid bleeding. One month after organoid transplantation, mice received 50 μl of 70 kDa Texas-Red dextran (Invitrogen, D1830) via retro-orbital intravenous injection. After 30 min, mice were euthanized and kidneys were collected. Explanted tissues were fixed and sliced for further immunostaining as described above.

**TUNEL staining**

Frozen sections were fixed with 4% PFA for 30 min and permeabilized with 0.2% Triton X-100 in pre-cold PBS for 10 min. The sections were then incubated with 50 μL of TUNEL reaction mixture (Vazyme Biotech, A113-02) at 37℃ for 60 min following the manufacturer’s protocol. After washing, samples were incubated with primary antibodies overnight at 4℃, followed by incubation with secondary antibodies for 1 h at room temperature. Nuclei were stained with Hoechst 33342 and the sections were imaged using a fluorescence microscope (Olympus, FV3000). The antibodies are listed in Table S2.

**Western blot**

Total proteins were extracted from organoids using RIPA lysis buffer (Solarbio, R0010) supplemented with protease inhibitor cocktail (MedChemExpress, HY-K0011). Protein concentration was quantified using the Pierce BCA Protein Assay Kit (Beyotime, p0012). Thirty micrograms of total proteins were resolved on 12.5% SDS-PAGE gel and transferred to a 0.4-μm polyvinylidene difluoride membranes. Membranes were blocked with 5% non-fat milk in TBST (100mM Tris–HCl pH 7.4, 0.9% NaCl, and 0.1% Tween 20) for 1 h at room temperature, followed by overnight incubation with primary antibodies at 4℃. After three washes in TBST, membranes were incubated with corresponding secondary antibodies, and protein bands were visualized using an BioRad imaging system. The antibodies used for western blot are listed in Table S2.

**Seahorse Mito Stress test**

Mitochondrial bioenergetic parameters were analyzed according to the Seahorse Mito Stress test kit protocol (Agilent). Organoids were plated in a Matrigel-coated 96-well XF plate (Agilent). The oxygen consumption rate (OCR; pmol$O_2$/min) measurements under basal condition and after the sequential injection of the ATP synthase inhibitor oligomycin A (2.5 μM), the ETC accelerator ionophore FCCP (Carbonyl cyanide-p-trifluoromethoxyphenylhydrazone, 1 μM), and the ETC inhibitor mixture of rotenone (2 μM) and antimycin A (2 μM). All data were normalized by protein concentration.

**EC isolation from hBVOs and vhCOs**

hBVOs or vhCOs were dissociated into single-cell suspension as described above. Once single cells were obtained, the digestion reaction was quenched using cold CDM medium supplemented with 10% KOSR. Cells were collected by centrifugation at 300x g for 5 min at room temperature and resuspended into MACS® buffer consisting of 0.5% MACS® BSA Stock Solution and 2 mM EDTA in DPBS. The cells were passed through a 40 μm filter to ensure that single cells were collected. CD144-expressing ECs were then isolated using the human CD144 (VE-Cadherin) MicroBeads and LS Columns (Miltenyi Biotec, 130042401) according to manufacturer's instruction. RNA was extracted and analyzed by qPCR as described above.

**Processing and analysis of 10x genomics single-nucleus multiomic library**

Nuclei were isolated following the 10X Genomics Nuclei Isolation protocol. 100,000 cells were lysed with 0.1% lysis buffer (10mM Tris-HCL pH 7.4, 10mM NaCl, 3mM MgCl2, 0.1% NP40, 1% BSA) for 3 min on ice. Lysis efficiency and nucleus quality were assessed by trypan blue staining. RNA-seq and ATAC-seq libraries were prepared with the 10X Genomics Chromium Next GEM Single Cell Multiome ATAC + Gene Expression Kit and sequenced as recommended. Quality control was conducted after cDNA amplification and ATAC library construction by Bioanalyzer analysis and Qubit measurement. Libraries were pooled and sequenced according to 10x Genomics recommendations on an Illumina NovaSeq X plus system.

RNA and ATAC reads from 10x Multiome single-nucleus RNA (snRNA) and ATAC sequencing were preprocessed, aligned, and quantified using Cell Ranger software package v 2.0.0 (10x Genomics) and human reference hg38 (10x Genomics refdata-gex-GRCh38-2020-A). Nuclei that passed Cell Ranger ARC filters were analyzed with Signac, filtering for RNA data with 200–10,000 genes and less than 15% mitochondrial reads, and ATAC data with at least 1,000 unique molecular identifiers, a nucleosome signal less than 2, and a TSS enrichment score greater than 1.In addition, DoubletFinder (v2.0.3) was applied to the RNA data to identify and remove potential doublets.

**Dimensionality reduction, integration, and clustering**

Downstream analysis was performed using Seurat v4.4.0 and Signac v1.11.0. For gene expression, counts were log-normalized, and the top 2,000 highly variable genes (HVGs) were scaled. The top 25 principal components (PCs) were calculated, and batch effects were corrected using Harmony. For chromatin accessibility, the peak matrix underwent TF-IDF normalization and latent semantic indexing (LSI), utilizing dimensions 2–30.

Multimodal integration was achieved using Weighted Nearest Neighbor (WNN) analysis to generate a joint UMAP visualization. Clusters were identified using the Louvain algorithm. Cell types were annotated using SingleR v1.4.1, followed by manual curation based on marker genes (CellMarker 2.0, PanglaoDB) and chromatin accessibility tracks visualized in CoverageBrowser. Subpopulations were identified by re-clustering specific lineages using lineage-specific features.

**Trajectory inference**

Differentiation potential was quantified using CytoTRACE 2 v1.0.0. RNA velocity analysis was conducted using scVelo v0.2.5. Spliced and unspliced matrices were filtered (min_shared_counts=20) and subset to the top 2,000 genes. The dynamical model was employed to estimate velocity vectors, which were projected onto the UMAP embedding.

#### Regulatory network and cell-cell communication analysis

Transcriptional regulatory networks were inferred using pySCENIC v0.12.1. Co-expression modules were generated using GRNBoost2, and motif enrichment was performed using the CisTarget database (hg38) to identify regulons. Regulon activity per cell was scored using AUCell.

Intercellular communication was analyzed using CellChat v2.1.2. Overexpressed ligand-receptor interactions were identified within cell types and aggregated to signaling pathways to evaluate differential communication strength and patterns.

#### Functional enrichment

Differentially expressed genes (DEGs) were analyzed for functional enrichment (GO, Reactome, WikiPathways) using clusterProfiler v3.18.1 (Benjamini-Hochberg adjusted P $< 0.05$). Global pathway activity scores were computed for single cells using GSVA v1.48.0. Differential pathway activity between conditions was assessed using linear modeling in limma v3.46.0.

#### Maturation assessment

To quantify cardiomyocyte maturation, a maturation index was calculated based on the expression ratios of developmental isoform pairs: *TNNI3/TNNI1* and *MYH7/MYH6*. A pseudocount of 0.01 was added to mitigate zero-inflation issues.

**Public data analysis**

The public datasets GSE202398, GSE134355, GSE156793, and GSE114607 used in this study were downloaded as raw data from the Gene Expression Omnibus (GEO) database. Human fetal cardiac tissue data at 6.5 weeks of gestation were obtained from the Spatial Research website (https://www.spatialresearch.org/resources-published-datasets/). All datasets were subsequently filtered and analyzed using the parameters established in their respective original studies.

**QUANTIFICATION AND STATISTICAL ANALYSIS**

Comparisons between two groups were performed using Student's t-test. While comparisons among multiple groups were performed with One-way ANOVA. All data were presented as the mean ± SEM. A p value of $<0.05$ was considered significant. *p<0.05, **p<0.01, ***p<0.001, ****p<0.0001. Statistical details can be found in the figure legends. n represents different specimens from different cell lines or biological replicates. Sample size for all experiments was specified in figure legends. Violin plots were plotted with Bonferroni correction test and FDR<0.05 was regarded as significant. Analyses were carried out using GraphPad Prism 10.0. As for all the experiments, at least three independent experiments were performed to reach the minimal requirement for statistical analysis, unless otherwise specified. Blinding or randomization was not performed unless otherwise specified. Exclusion is not applied in this study.
