## Supplementary figures and images for "Vascularized Human Cardiac Organoids Reveal Endothelial–Cardiomyocyte Crosstalk and Mechanisms of Carfilzomib-Induced Cardiotoxicity"

### Supplementary figures

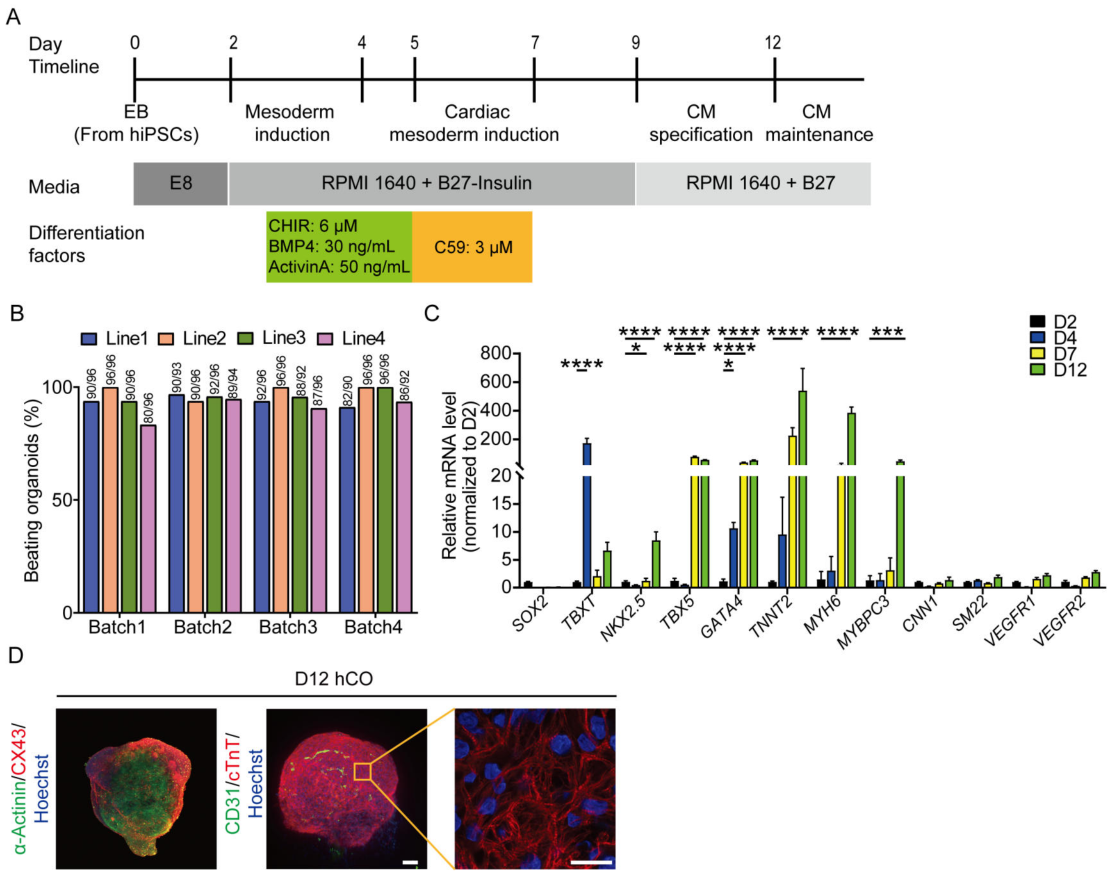

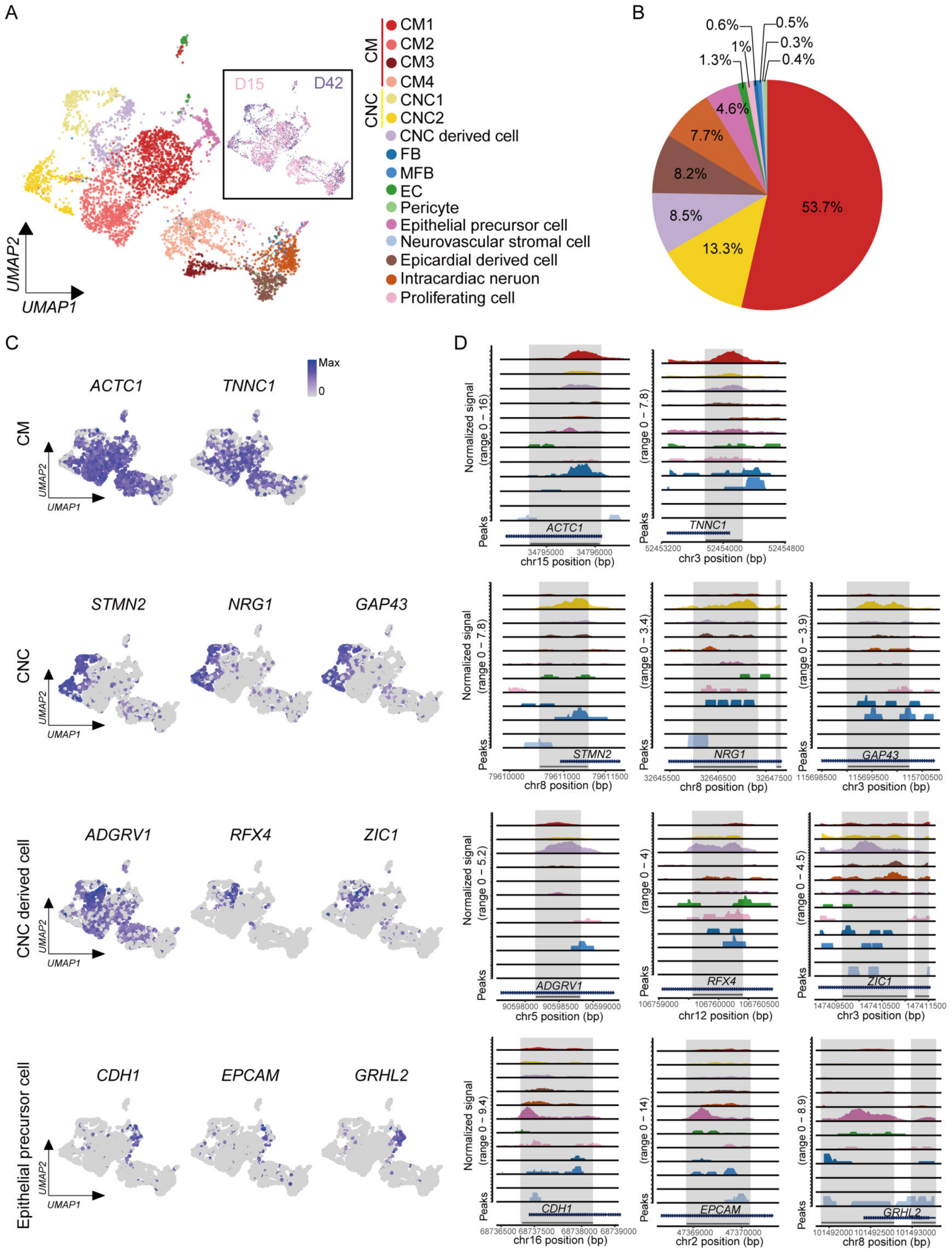

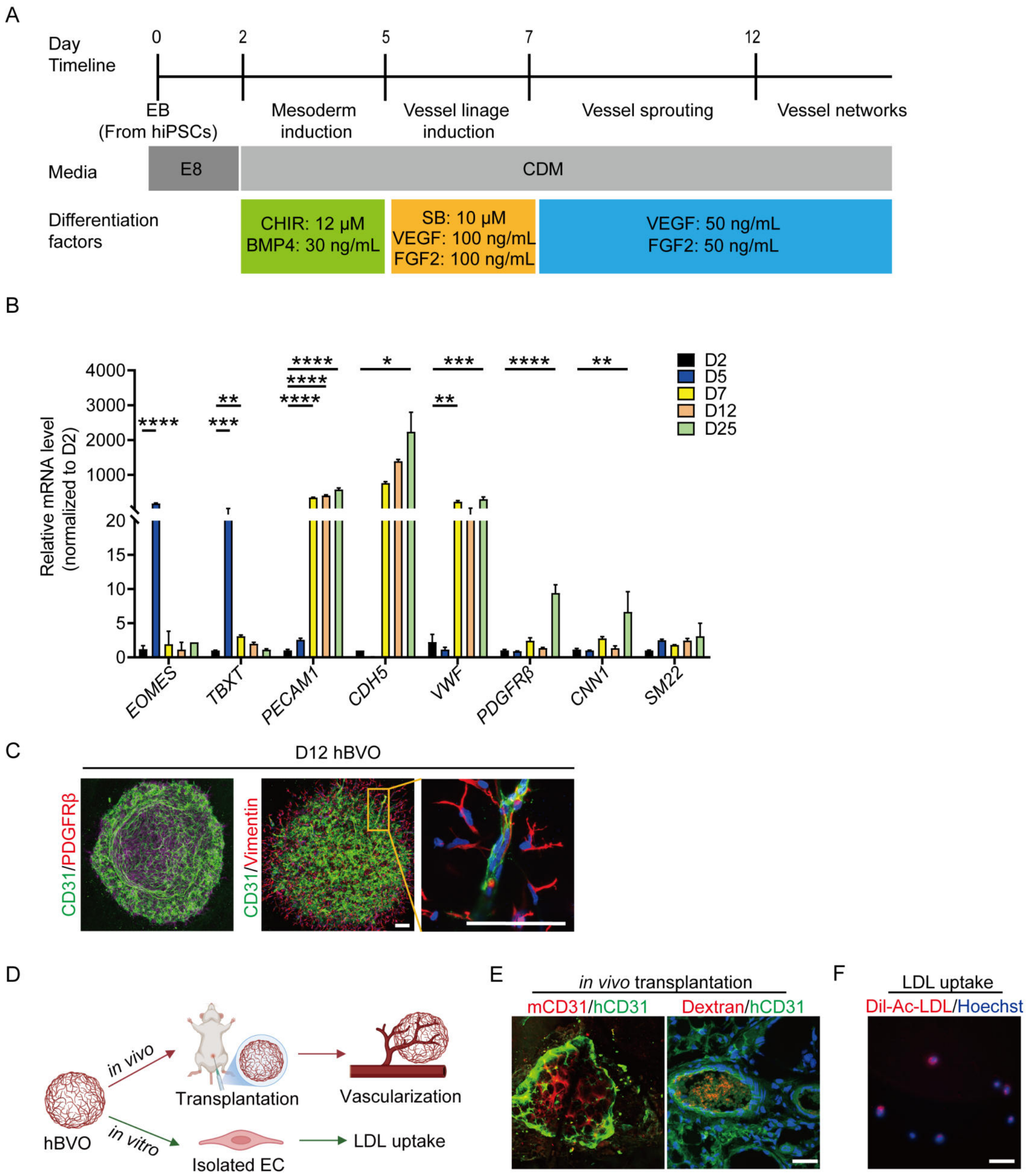

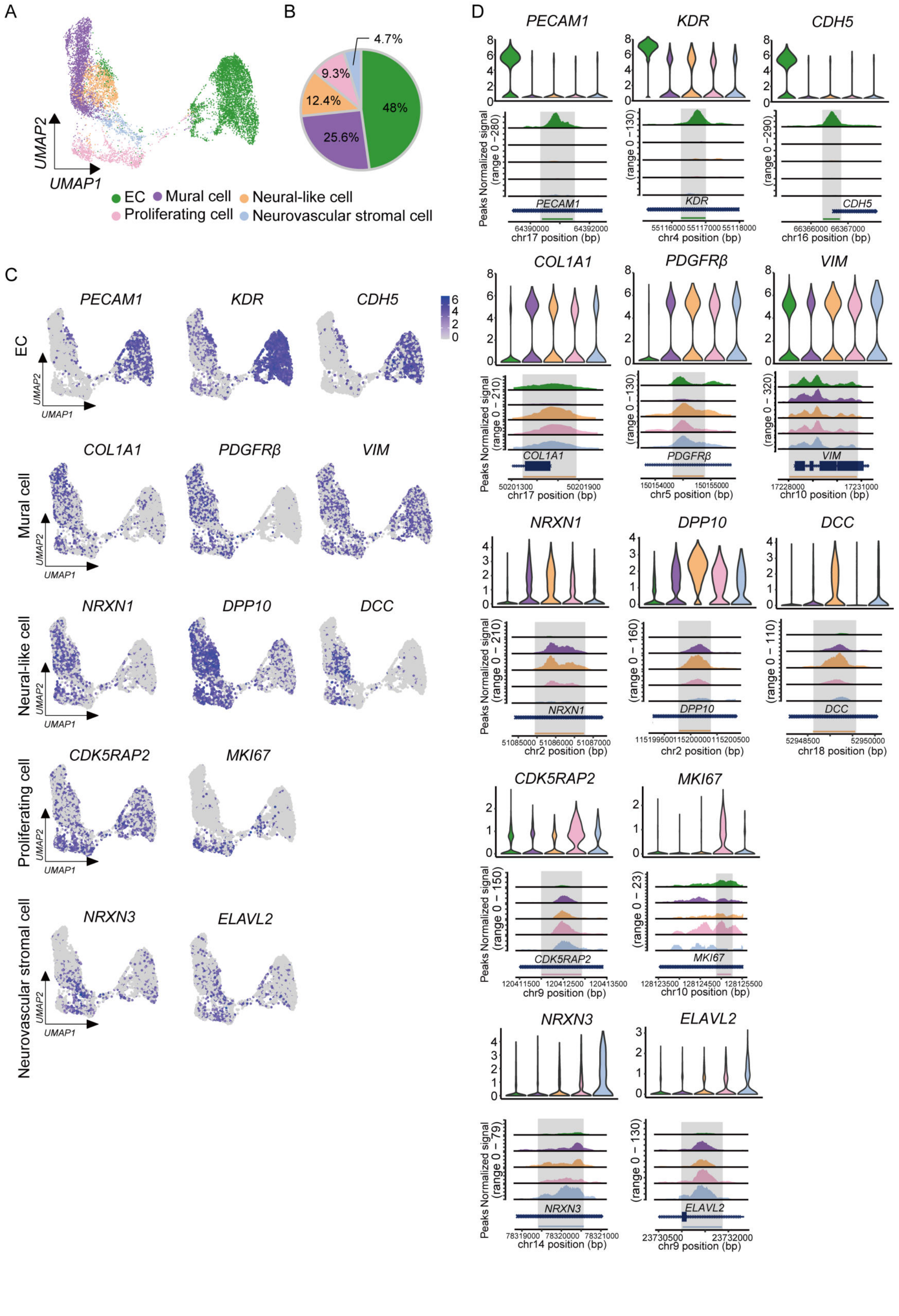

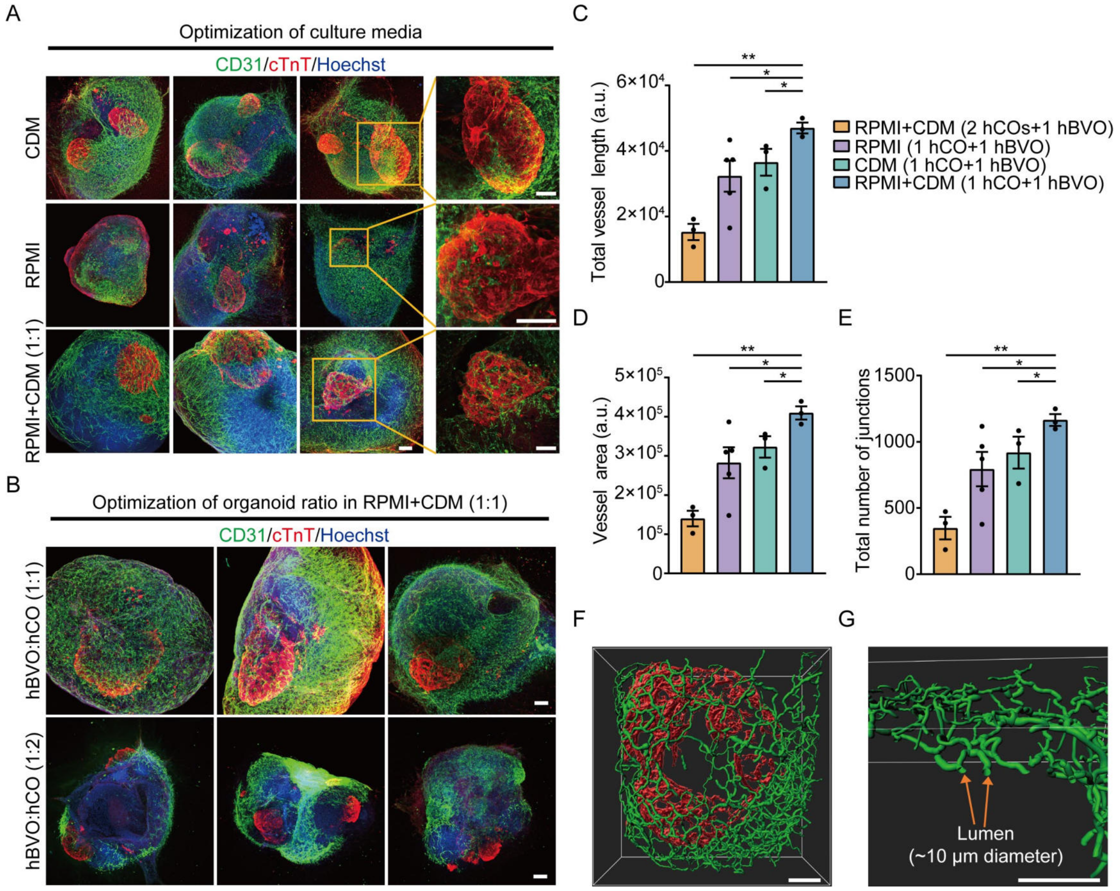

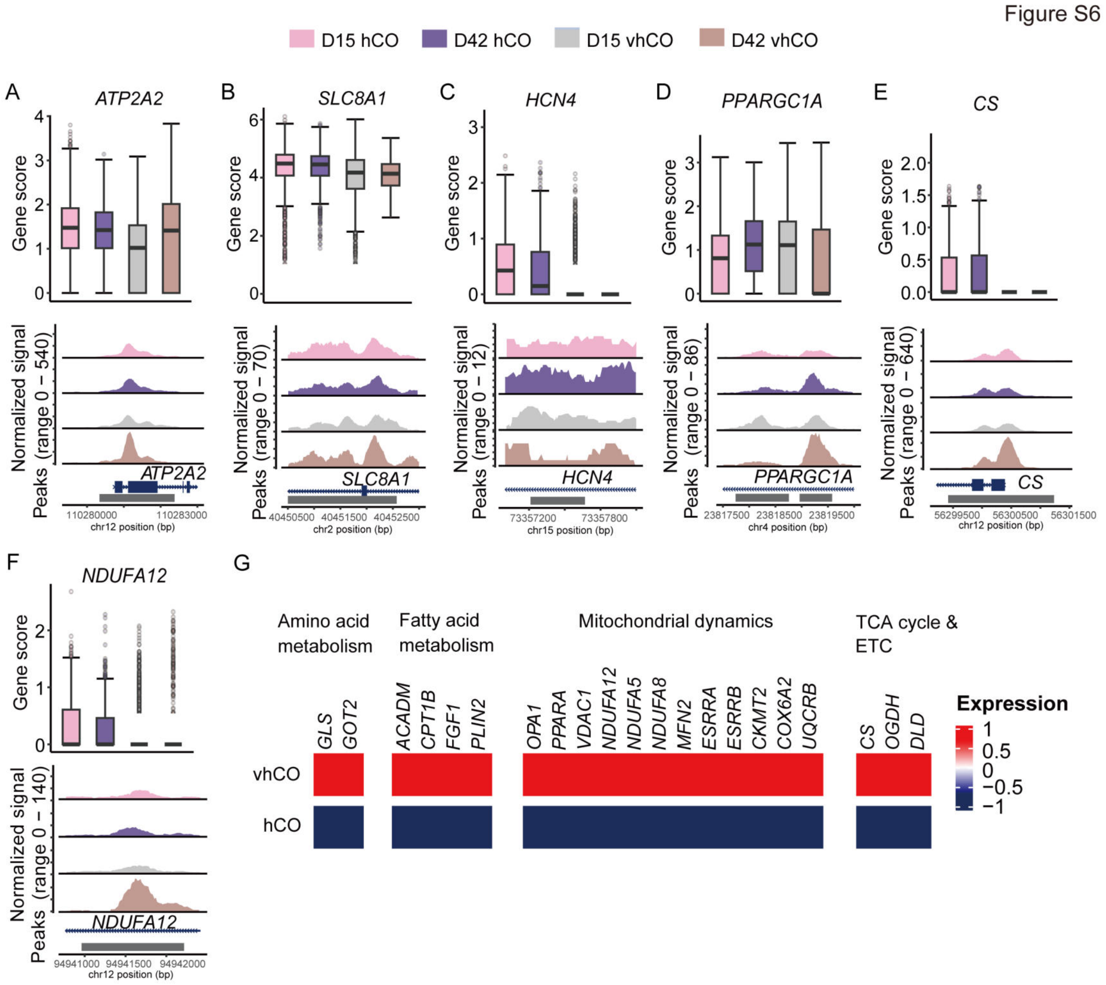
